# BiGER: Bayesian Rank Aggregation in Genomics with Extended Ranking Schemes

**DOI:** 10.1101/2025.06.17.660022

**Authors:** Kaiwen Wang, Yuqiu Yang, Yusen Xia, Guanghua Xiao, Johan Lim, Xinlei Wang

**Affiliations:** Department of Statistics and Data Science, Southern Methodist University, Dallas, TX, USA, 75275; Department of Public Health, Peter O’Donnell Jr. School of Public Health, University of Texas Southwestern Medical Center, Dallas, TX, USA, 75390; Robinson College of Business, Georgia State University, Atlanta, GA 30303; Quantitative Biomedical Research Center, Peter O’Donnell Jr. School of Public Health, UT Southwestern Medical Center, Dallas, TX, USA, 75390; Simmons Comprehensive Cancer Center, UT Southwestern Medical Center, Dallas, Texas; Department of Bioinformatics, University of Texas Southwestern Medical Center, Dallas, TX, USA, 75390; Department of Statistics, Soul National University, Seoul, Korea; Department of Mathematics, University of Texas at Arlington, Arlington, USA, 76019; Division of Data Science, College of Science, University of Texas at Arlington, Arlington, USA, 76019

**Keywords:** Bayesian hierarchical modeling, gene expression, Gibbs sampling, meta-analysis, posterior inference, variational inference

## Abstract

With the rise of large-scale genomic studies, large gene lists targeting important diseases are increasingly common. While evaluating each study individually gives valuable insights on specific samples and study designs, the wealth of available evidence in the literature calls for robust and efficient meta-analytic methods. Crucially, the diverse assumptions and experimental protocols underlying different studies require a flexible but rigorous method for aggregation. To address these issues, we propose BiGER, a fast and accurate Bayesian rank aggregation method for the inference of latent global rankings. Unlike existing methods in the field, BiGER accommodates mixed gene lists with top-ranked and top-unranked genes as well as bottom-tied and missing genes, by design. Using a Bayesian hierarchical framework combined with variational inference, BiGER efficiently aggregates large-scale gene lists with high accuracy, while providing valuable insights into source-specific reliability for researchers. Through both simulated and real datasets, we show that BiGER is a useful tool for reliable meta-analysis in genomic studies.

## INTRODUCTION

Recent years have seen the rise of high-throughput assays in addressing important biological questions, such as disease phenotyping^1^, differential expression^2^, and downstream modeling^3^. This is especially true for single-cell RNA sequencing (scRNA-seq)^4^ experiments in which differential expression analysis forms part of the core analysis workflow. It is increasingly common that researchers utilize such tools to report significant genes, whether differentially expressed or otherwise experimentally validated, across conditions and diseases of interest. When examining results on the same general topic (*e*.*g*. non-small-cell lung cancer^5,6^ or COVID-19^7,8^), one is often faced with a cacophony of genes and gene lists with varying degrees of quality, underlying assumptions, and data formats. The robust aggregation of available information thus becomes an important undertaking during the process of meta-analysis.

Gene lists from differential expression analyses are commonly presented as ranked lists, often based on metrics like fold changes between conditions and p-values from the statistical tests used (*e*.*g*. Wilcoxon Rank Sum test in Seurat^9^). Within this framework, the problem of combining multiple gene lists becomes that of rank aggregation (RA), a topic extensively explored in bioinformatics^10,11^ and other computational fields^12^. A key idea behind using RA in meta-analysis is that an aggregated gene list can provide a more reliable consensus ordering than any single list on its own. However, while the underlying principle of RA seems straightforward, a number of complications exist regarding the characteristics of the gene lists themselves and decisions about which studies to include.

What is often overlooked is the heterogeneity of reported gene lists in the scientific literature. More specifically, there are three unique scenarios that are nontrivial for existing RA methods. Firstly, it is often the case that the exact rankings are not specified, occurring when there are actual ties or when the ordering is not given in the publication. This phenomenon can result in many possible ranking schemes, and it complicates the modeling process by involving ties and possible tie-breaking procedures^13,14^. Secondly, one cannot assume that all gene lists are complete. Namely, studies may either neglect to report non-significant genes or fail to assay a subset of genes due to technical reasons. Thirdly, the process of meta-analysis induces what is so-called researchers’ degree of freedom, where the selection of studies is crucial but the procedure of which is not always clear^15^. Under the modus operandi of letting data speak, the ability to discern study relevance and quality is instrumental for an RA method to succeed and for researchers to understand the results. These complications in the RA of gene lists entail the design of methods that can handle complex input data formats with study quality assurance in the estimation procedure.

Previously, many RA methods have been proposed and adopted for the purpose of meta-analysis. In fact, two comprehensive review studies^5,16^ observed a few important trends on the existing methodologies in the field. From a most simplistic standpoint, Borda’s collection^17^ with simple statistics serves as a sometimes adequate but often *ad-hoc* baseline for analyses. Many Bayesian methods are statistically more robust by modeling the underlying data-generating mechanisms of gene lists. Specifically, BIRRA^18^ combines traditional Borda’s method with Bayes factor to produce an aggregated ranking; BARD^19^ and BiG^6^ are both Bayesian hierarchical models implemented with Markov Chain Monte Carlo (MCMC)^20–22^, where the former models each item’s relevance and the latter utilizes a latent variable approach with support for unreported genes. An improved version of BARD with capability for estimating ranker quality using Mallows model was later proposed^23^. A common issue with Bayesian methods, especially complex models with MCMC, is the lack of scalability in genomics with a growing body of literature. On the other hand, there also exists many frequentist methods in the field of RA and bioinformatics. For example, Stuart’s method^24^ is developed for detecting co-expression of genes in DNA microarrays using order statistics, whereas Robust rank aggregation (RRA)^25^ is based on hypothesis testing by comparing rankings to a null model. More recently, MAIC^8,26^ innovatively incorporated studies in which reported genes are not specifically ranked, significantly broadening the possible scope of meta-analysis in the context of RA.

Advancements in RA highlight the critical need for approaches that are broadly applicable, inherently interpretable, and computationally efficient. Although existing methods often perform well in certain aspects, they typically struggle to achieve high performance across all three simultaneously. To address this challenge, we introduce **B**ayesian Rank Aggregation **i**n **G**enomics with **E**xtended **R**anking Schemes (BiGER), a statistical technique specifically designed to achieve robust and efficient RA while also incorporating interpretable posterior estimations on gene ranking and study quality. BiGER leverages a flexible Bayesian hierarchical model, coupled with variational inference (VI) ^27,28^, to effectively handle large-scale input lists with diverse ranking schemes for efficiency and generalizability.. Extensive validation on three established benchmark datasets^16^ and 25 in-house simulation scenarios show that BiGER consistently ranks among the best-performing methods in accurately identifying significant genes, irrespective of input list type. BiGER also demonstrates resilience against the inclusion of irrelevant studies in meta-analyses, ensuring reliable performance.

The generalizability of BiGER arises from its ability to handle all possible types of gene lists that occur in genomic studies. Specifically, it provides formal treatment to reported vs. unreported and ranked vs. unranked genes in each contributing study. By contract, BiG^6^ cannot handle top unranked gene lists and MAIC cannot handle mixed lists that contain both top ranked and top unranked genes. Explainability is inherent in its Bayesian foundation and explicit modeling with directly interpretable parameters, providing insights into reliability of individual studies. Efficiency is achieved through its VI-based computational approach, suitable for large genomic meta-analyses. Built on BiGER that effectively balances generalizability, interpretability, and efficiency, we thus present a rigorous while easy-to-use pipeline as our deliverable for researchers to conduct future meta-analyses across domains.

## RESULTS

### Defining ranking schemes in genomic studies

The BiGER workflow consists of researchers first collecting a set of relevant studies from the literature, then obtaining gene lists, and subsequently performing RA using the method we propose (**Fig. 1a**). To define the scope of BiGER and its relevance in the RA literature, we start by introducing the overall framework with notations and assumptions. Suppose there are □ genes of interest in total indexed by □ and □ selected studies indexed by □. We let 𝒢 denote the full set of genes observed across all studies, with |𝒢| = *G*. Given that each study may not report or study all the genes in 𝒢, the gene set for each study □ satisfies the relationship 𝒢_*j*_ ⊆ 𝒢. A full list of notations and parameters are given in **Supplementary Note 1**.

**Fig. 1.**
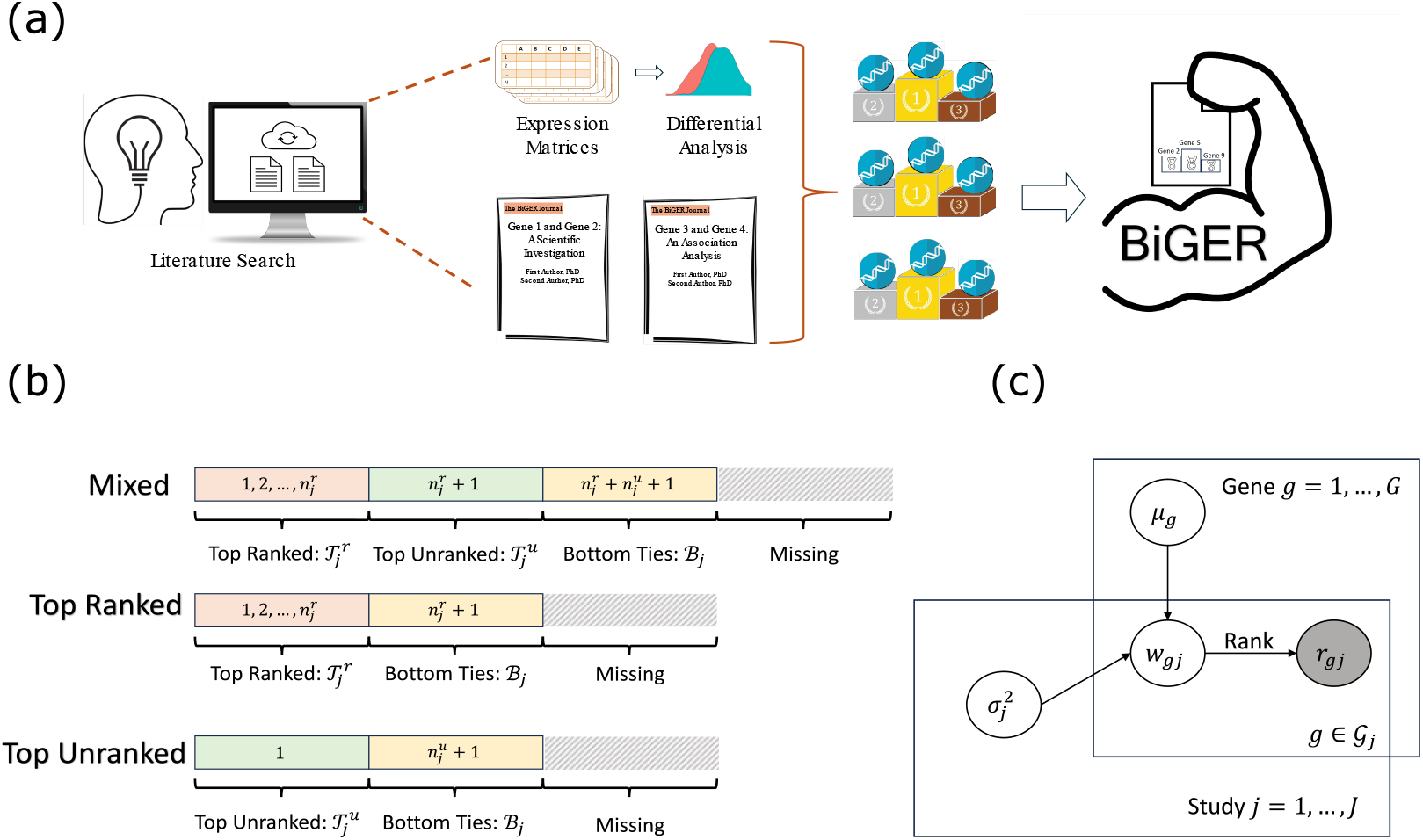
The overview of the framework of BiGER and its statistical foundation. (a) A visual pipeline for a typical meta-analysis pipeline. (b) Structures of gene lists supported in BiGER. Each block represents types of genes in a given study with its associated with ranks. (c) A plate diagram of the BiGER model. The filled grey circle represents observed ranks in a meta-analysis.

Gene lists observed in genomics literature often exhibit commonalities that are more complex than full ranked lists. In BiGER, we distinguish between four types of genes encountered in meta-analysis (**Fig. 1b**). First, one of the most common is top-ranked genes, whose ranks start from 1 (the most important) and are explicitly given. We let 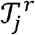 denote the set of the top-ranked genes in study *j*. For notation in general, we use letter superscripts for index purposes instead of exponents unless otherwise noted. Second, when the exact ranks are not given but a gene list is still reported, we assume such genes to be tied to the best of our knowledge. They are thus called top-unranked genes from the set 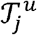. The third and fourth categories deal with unreported genes for each study because it is rare that a study comprehensively reports all genes involved in a meta-analysis. Following the convention of BiG^5^, if a study investigated some genes but failed to report them as significant, then they are assumed to be bottom ties. As the name suggests, bottom ties are categorically less important than top-ranked and top-unranked genes. We denote this set as ℬ_*j*_. On the other hand, when a study simply did not analyze the missing genes due to poor-quality or no data, then we treat them as missing data (NA). In this case, we resort to all-available case analysis without making any additional assumption on the missingness.

To allow for flexibility in meta-analysis and account for unique studies, we allow studies to have a combination of the four types of genes with some assumptions. Specifically, we define studies based on top genes in each study. If a study has only top-ranked genes without top-unranked genes (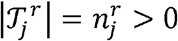 and 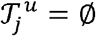), it is defined as a top-ranked study. Conversely, a study with top-unranked genes but without top-ranked genes is a top-unranked study 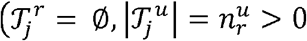, and 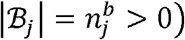. Given that the definitions of top-ranked genes and top-unranked genes are not mutually exclusive, it is possible that a study can report both types of genes. For example, a study may identify a short list of important genes, which are subsequently validated via experiments or downstream analyses. At the same time, the same study may report a list of less important but still relevant genes for completeness. Such studies are thus known as mixed studies with a mixed gene list 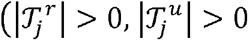 and 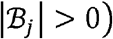. It is natural to assume that top-ranked genes are more significant than top-unranked genes in a mixed study. This assumption is important to distinguish between top-unranked genes and bottom ties: the former is explicitly reported by the study, and the latter needs to be assumed/inferred when performing RA.

In practice, any of the three types of studies may contain bottom-tied and missing genes. Further, a meta-analysis can easily contain many types of studies, each with a different configuration from **Fig. 1b**. A more in-depth discussion of some special cases is included in the **Methods** section.

### Overview of statistical framework

As a foundation of our model, we assume that genes in each study are ranked using some measure. In some cases, such information may be reported, such as fold change from differential analysis, but for generality, we assume that that the underlying scale is latent. Let ***ω*** be a *G* × *J* matrix whose element *ω*_*gj*_ denotes the latent weight measuring the local importance of gene *g* in study *j*, ***µ*** be a *G* × *J* vector with element *µ*_*g*_ reflecting the global importance of gene *g*, and ***ϵ*** be a *G* × *J* matrix with element *ϵ*_*g j*_ denoting the random error arising from study *j* for gene *g*. Then we can posit the following linear additive model

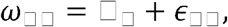

where we assume *µ*_□_ ~ □ (*0,1*) to ensure identifiability and 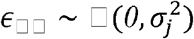, where the study-specific precision 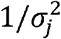 represents the overall quality of study *j* and *N* (·) is a normal distribution parametrized by the mean and variance parameters. Under these assumptions, the latent weight for each gene in a given study has the distribution 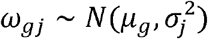.

Under BiGER’s framework and common RA data structures, the observed data consists of ranks of genes in each study, which we denote as ***r*** overall. In our latent variable approach, the order of latent weights **ω**_*j*_ for a given study *j* must be concordant with the observed ranking ***r***_*j*_; in other words, the observed *r*_*j*_ is determined by ***ω*** _*j*_. Therefore, the likelihood is an indicator function of the following form *p* (***r***_*j*_ ***ω***_*j*_) *= I* (The order of ***ω***_*j*_ is correct), where it is 1 if the latent weights are ordered correctly and 0 otherwise. While this is conceptually simple, we provide a complete form of *p* (***r***_*j*_ |***ω***_*j*_) in the **Methods** section with rigorous mathematical definitions. As part of our Bayesian framework (**Fig. 1c**), we further employ a conjugate prior *IG* (*a,b*) for 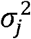’s, where *IG* (·) represents the Inverse-Gamma distribution using the shape-rate parameterization; *a* and *b* are hyperparameters, the choices of which will be further discussed. Using the notation **Θ** to represent the collection of all parameters and latent variables in our model, the full probability model is therefore written as

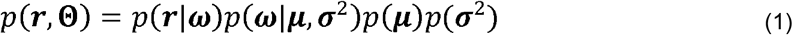

Posterior sampling and inference of ***µ*** and ***σ***^2^ can be achieved via a Gibbs sampler^20–22^. Details of the Gibbs sampler, hyperparameter selection, and software implementation are further discussed in the **Methods** section. We term this implementation the gBiGER method (Gibbs BiGER). As it represents the full model with asymptotically exact sampling, gBiGER serves a reference line for approximation methods aimed at faster posterior inference for large gene lists.

### Variational inference enables gene list aggregation at scale

While Gibbs sampling for gBiGER is adequately efficient for many applications as compared to BiG, large-scale meta-analysis of gene lists require even further optimization. In particular, variational inference (VI)^27,28^, a class of Bayesian approximation techniques, is a popular choice for conjugate models. Using the framework of mean-field VI (MFVI)^29^, we posit a family of independent variational distributions to approximate the posterior distribution of BiGER as

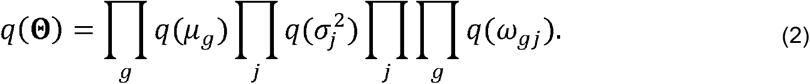

Following the convention of VI, we use the notation of *q* (·) to denote variational distributions, which is distinguished from the likelihood components. The optimization of such variational distributions is derived by minimizing the Kullback-Leibler (KL) divergence between the joint posterior distribution *p* (**Θ**|***r***) and *q*(**Θ**), which in turn is equivalent to maximizing the evidence lower bound (ELBO). It has been shown^29^ that the optimization of the target function ELBO results in the equivalence of finding

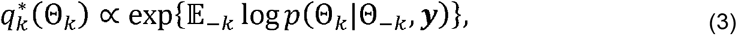

where we use a generic subscript *k* to index the variational parameter of interest, and subscript − *k* indexes all other parameters except for the *k*-th. The full conditionals used here are analogous to those from a Gibbs sampler.

While the conventional formulation of MFVI works well for most conjugate models, the likelihood of BiGER poses an issue for the derivation of the variational family. Namely, we previously defined the likelihood for a list to be *p* (***r***_*j*_ │***ω***_*j*_) = *I* (The order of ***ω***_*j*_ is correct). Without loss of generality, suppose we consider an arbitrary gene, which is situated in the middle of the top-ranked portion of the gene list. Its likelihood contribution is written as 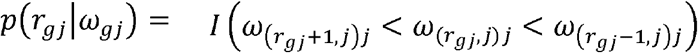 in which the first part of the subscript (*r*_*gj*_, *j*) returns the index of the item that is ranked *r*_*gj*_ in study *j*. It is obvious that the latent weight *ω*_*gj*_ depends on its neighbors (see the **Methods** section for a general treatment): while this poses no issue for a Gibbs sampler with full conditional distributions, expectations in MFVI will not have closed-formed solution under this formulation due to the need for computing expectations (see **Supplementary Note 1**). We thus propose an approximation scheme to solve this problem.

The approximation is conceptually simple: define an approximation via fixing the upper and lower bound for *ω*_*gj*_. This approach is justifiable because the overall BiGER framework cares only about the order of ***ω***_*j*_ via an indicator function, and the exact value of each latent weight has no further constraint. Further, it is implicit in the BiGER model that what is so-called data is ***r***_*j*_, which are observed ranks and depend only on the order of ***ω***_*j*_. For each study, we thus define a 1-1 transformation of *r*_*j*_ to formulate the boundaries of ***ω***_*j*_, which will be used in place of observed data (**Methods** section). Let 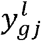 and 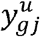 be the lower and upper bounds for *ω*_*gj*_ respectively, and the likelihood can be rewritten as 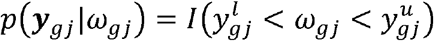. This parametrization treats ***y***_*gj*_ as data, which is fully observed, thus resolving the issue of dependence between parameters.

Incorporating the boundary-approximation method, we derive namesake BiGER algorithm using the Coordinate-Ascent VI (CAVI) algorithm as shown in Blei et al.^29^. We use the variational family previously specified to approximate the following joint posterior:

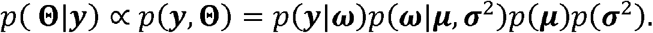

All variational distributions can be found in closed forms under this setting with parametrization similar to that of a Gibbs sampler in gBiGER. The details of the variational updates, the CAVI algorithm, and their corresponding derivations are included in the **Methods** section and **Supplementary Note 1**. In the rest of this work, BiGER is referred to as the VI-based implementation, whereas gBiGER is referred to as the Gibbs-based implementation.

### BiGER accurately models the global ranking of genes

To validate the performance of BiGER with known ground truth, we devised a comprehensive simulation scheme (**Methods** section) by varying many key settings one-at-a-time. Each simulated cohort is designed to consist of mixed studies as well as top-ranked and top-unranked studies to reflect the complex nature of a real meta-analysis. The exact type of each study is determined probabilistically using a categorical distribution. On average, each cohort is expected to have 20% mixed lists, 30% unranked lists, and 50% ranked lists. We repeat each simulation scenario 25 times to ensure results are repeatable. The details of the simulation are included in the **Methods** section.

In practice, there exist important factors that can affect the validity and practicality of any chosen RA method in meta-analysis. Specifically, we designed our simulation procedure to systematically investigate commonly encountered scenarios. Namely, we varied the number of studies (# Studies), number of genes (# Genes), expected proportion of missing genes (Expected % NAs), and overall study quality. The number of studies directly correlates with the amount of evidence available in a given field, but at the same time, it also reflects the potential consequences of conducting meta-analysis with insufficient literature search. Given the diverse applications of BiGER for the meta-analysis of many diseases, the number of genes and the quality of each study can vary significantly from application to application. We thus specifically simulate a varying range of correlations between true global importance and observed ranks for heterogeneous cohorts of studies, which mimic the reality.

We compared the performance of the BiGER family to BiG, MAIC, and methods from the “RobustRankAggreg” R package^25^, including the Borda’s collection, RRA, and Stuart’s method^24^. First, we investigated the coverage of each RA method across all the simulation settings. In general, coverage is defined as the percentage of truly significant genes (the truth set) covered in the top *K*_TOP_ genes produced by a given RA method. We set *K*_TOP_ = 100 unless otherwise noted. For simulation, we defined the truth set to be the top 10% of the gene set. To compare the coverage across different settings (**Fig. 2a**), we ranked each method within a given setting (column-wise in the plot). As a result, both BiGER and gBiGER are uniformly the top two methods across the board, which are shown with the darkest orange colors. There is no obvious coverage difference between BiGER and gBiGER with them trading the top spot depending on the configuration. The lack of performance difference between them provides us with good evidence for BiGER’s effective VI approximation. On the other hand, BiG, geometric mean, and Stuart’s method are also respectable performers, but they trail behind the BiGER family. MAIC inexplicably performs well in only some settings. The raw coverage values are presented as the dot sizes in the panel, and it is obvious that coverage correlates with some simulation settings, such as the number of genes and studies, prompting us to more closely inspect the effect of these settings.

**Fig. 2.**
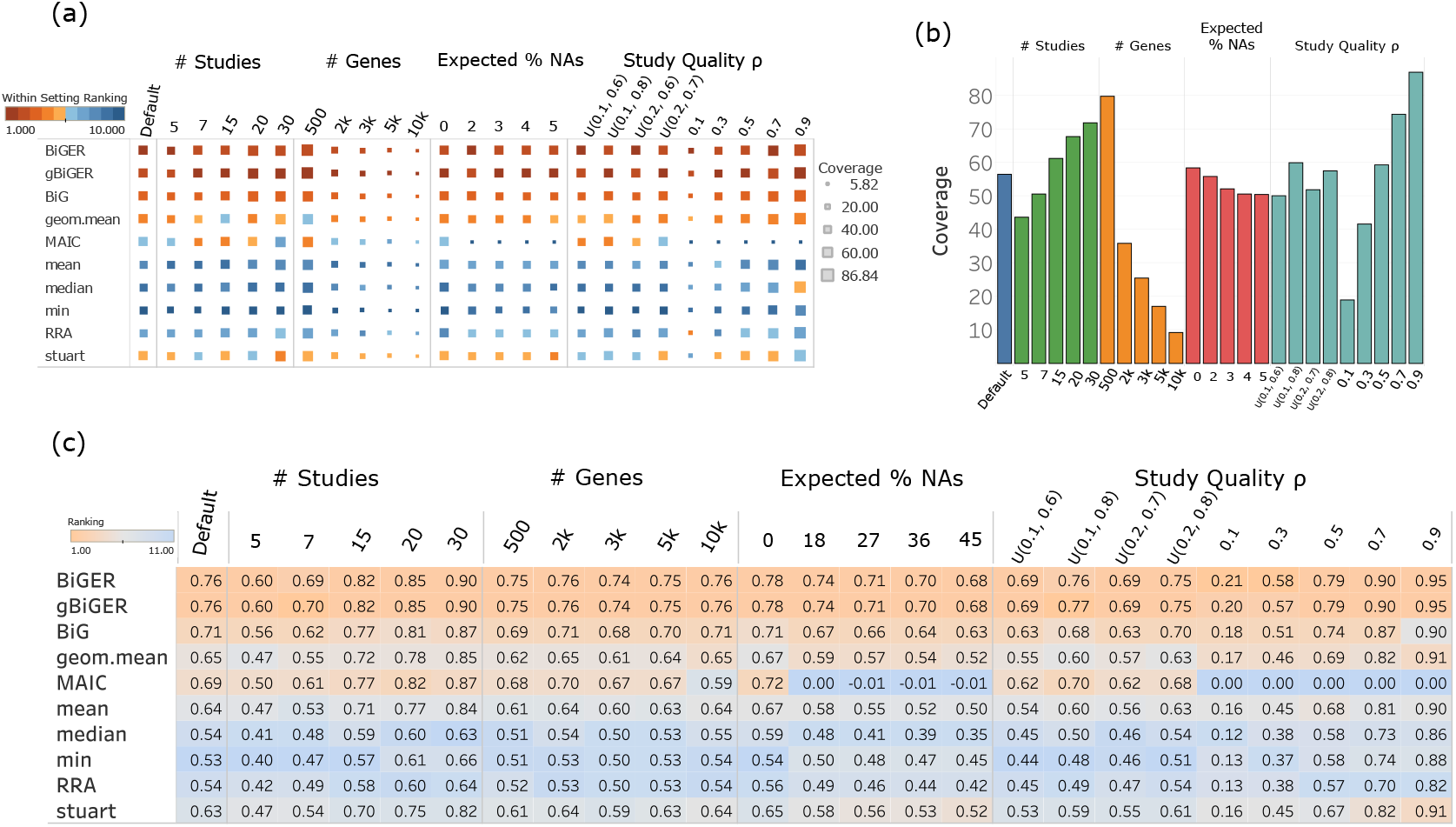
Accuracy validation of BiGER via comprehensive simulation. (a) A comparison of RA methods across all simulation settings. The metric is based on coverage. Both color and size of the squares are computed based on the top method of the category. Orange color and larger squares represent closer performance to the top performance, which has better coverage, and vice versa. (b) A bar plot as a summary of BiGER’s coverage across all runs under each setting. The metric used is coverage. Different colors denote the overall setting as denoted on top of the plot. (c) A table of spearman correlation to assess the performance of all methods in simulation. Colors within the table are based on the ranking of each method under the specific setting (column). Orange color denotes superior performance with larger spearman correlation, and blue colors denotes worse (smaller) spearman correlation.

Having established BiGER’s advantage over other methods, we shift our focus towards simulation settings by specifically plotting BiGER’s coverage in a bar chart (**Fig. 2b**). As expected, a few previously observed trends are further accentuated here. First, BiGER benefits from the inclusion of more studies, corresponding to an increasing sample size. On the other hand, the size of the gene set negatively impacts the overall coverage, due to the fact that more unrelated or marginally important genes are included. As designed, one key advantage of BiGER is its robustness against missing genes, as the coverage does not significantly deviate from the default despite the loss of information from simulated missing data. The same trend can be observed for cohorts with different overall study quality, where we use *ρ* to indicate the correlation between ***μ*** and ***ω***_*j*_ (see the **Methods** section for the full definition). When *ρ* is fixed, representing a meta-analysis of similar-quality studies, BiGER’s coverage directly reflects the study quality. In contrast, when *ρ* is sampled from a uniform distribution with a large range, BiGER’s posterior inference yields good coverage regardless. This robustness likely stems from BiGER’s ability to extract information from the sufficient-quality component studies. This latter scenario mirrors the heterogeneity often encountered in real-world literature.

Another key insight from RA is that of the exact order of posterior gene rankings. While coverage operates on the level of gene sets, the exact order is also valuable for practitioners who wish to investigate the relative importance of each gene individually. We thus computed the spearman’s correlation between the true ranking and the inference-based ranking for all methods across all settings (**Fig. 2c**). BiGER variants achieved comparably the best results across the board. Again, BiGER’s performance does not significantly differ from that of gBiGER. While we again observed the same trends as the coverage-based results in **Fig. 2a**, the pattern here is even more obvious. The BiGER variants form the first tier with BiG following with noticeable performance loss. MAIC, geometric mean, arithmetic mean, and Stuart’s method form the next tier, but MAIC produces nonsensical results under certain configurations with large ratios of missing data and with fixed study qualities. The remaining methods form the last tier with the lowest spearman’s correlation.

Comparing BiGER with the original BiG in both **Fig. 2a** and **Fig. 2c**, BiGER performs better across the board. As previously discussed, a real meta-analysis in genomics may consist of both mixed studies and top-unranked studies in addition to top-ranked studies. BiGER is the only method that supports these studies natively, which explains its superior performance and overall flexibility in accommodating different possible scenarios. While BiG is also a Bayesian hierarchical model capable of modeling bottom-tied and missing genes, it cannot model top-unranked genes from mixed or top-unranked studies. In this case, such studies have to be either discarded as in our benchmark or modified using *ad-hoc* procedures. BiG still outperforms many statistics-based methods (e.g., geometric mean) that also lack support for mixed or top-unranked lists, likely due to its rigorous Bayesian treatment of bottom ties and NAs. Despite this limitation, BiG still outperforms many statistics-based methods (*e*.*g*. geometric mean) that also lack support for mixed or top-unranked lists, likely due to its rigorous Bayesian treatment of bottom ties and NAs. Building upon this statistical foundation, BiGER generalizes BiG to achieve even better performance and more accurately model the observed data. The simulation study thus showcases BiGER advantage in modeling a wide range of conditions with many types of input lists.

### Rank Order Stability and Partial Gene Set Robustness

Despite the ease of literature search in the digital age, the comprehensiveness of a meta-analysis depends largely on the researcher. It is foreseeable that relevant studies may be left out due to various valid reasons. We therefore use our default simulation scenario to test how leaving out studies can affect the coverage of BiGER and other methods. Specifically, we selected the first default dataset (10 studies and 1,000 genes) and subsampled 3, 5, or 7 studies for computation. To ensure that the results were not affected by the specific seed chosen, we repeated the subsampling procedure 10 times for each setting.

Under the reference setting with all 10 studies, the BiGER family achieves the highest coverage across the board (**Fig. 3a**), again affirming the overall results seen in **Fig. 2a**. Expectedly, as we expand the *K*_TOP_ genes considered across the *x*-axis, the coverage monotonically increases until it approaches 100%. The performance is a bit mixed for other methods with BiG being near the top of the pack. When considering subsampling, a few striking patterns emerge. First of all, regardless of the number of studies sampled, the BiGER family consistently has the best coverage among all the methods. This phenomenon showcases a good rank-order stability of BiGER among all the RA methods. Specifically, BiGER is robust against selecting only a small subset of the studies, meaning that the benchmark previously done is still valid regardless of users’ thoroughness in literature search.

**Fig. 3.**
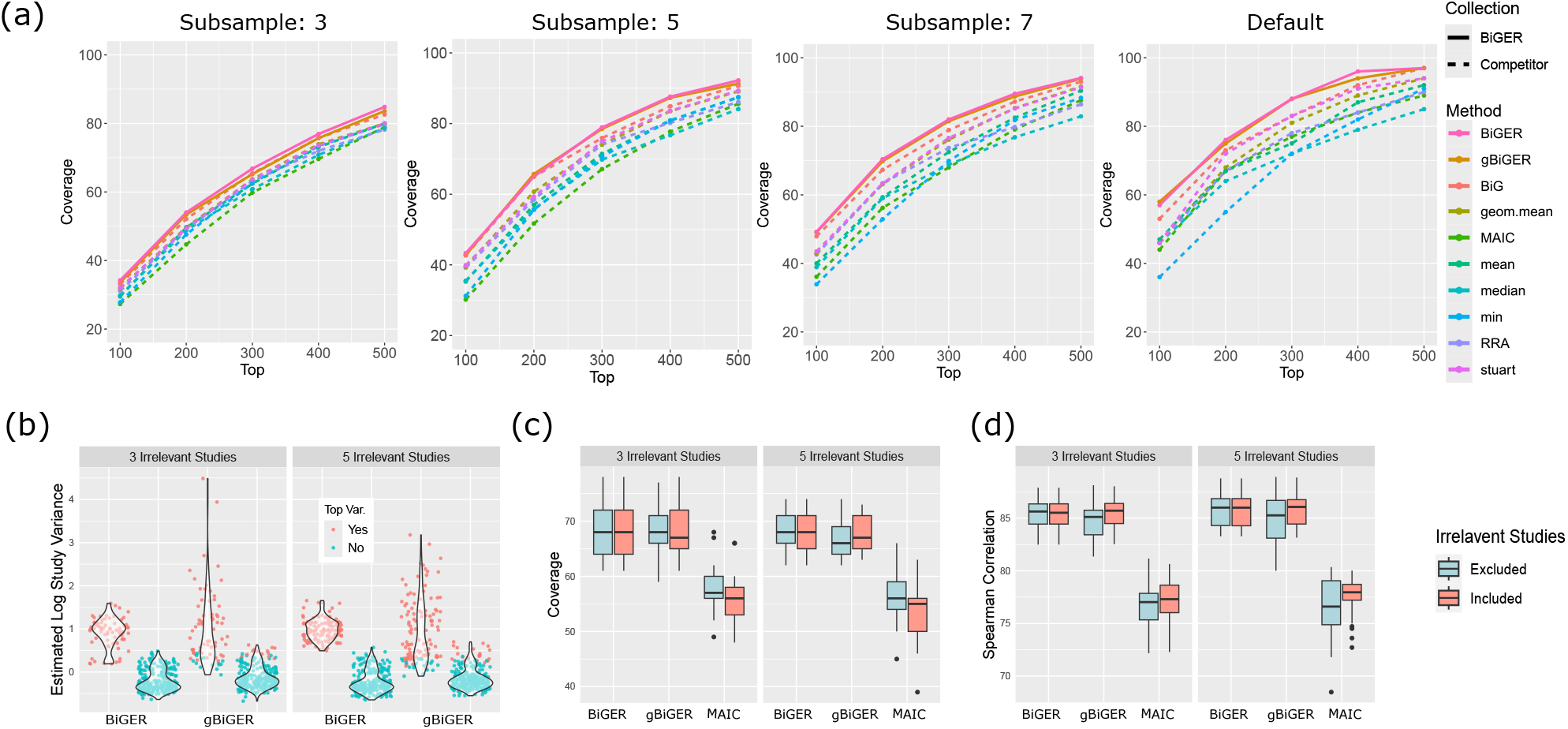
Robustness of BiGER in the presence of small cohort and irrelevant studies. (a) Line plots of the coverage of RA methods with subsampling. Each line with a different color represents an RA method. Solid lines denote the BiGER family of methods, whereas dashed denote all other methods. (b) Side-by-side violin plots of estimated log posterior variance with the inclusion of irrelevant studies. Each dot represents an individual study with all runs combined into a single violin for each method. Red color indicates that the study is ranked in the top 3 or 5 in terms of posterior variance within the corresponding pane. Blue color indicates the rest of the studies. The left-side violin for each method consists of irrelevant studies, and right-side violin consists of informative studies. (c) A side-by-side comparison of the effect of including irrelevant studies using coverage. Red boxes indicate runs with irrelevant studies included, whereas teal boxes with irrelevant studies removed. (d) A side-by-side comparison of the effect of including irrelevant studies using spearman correlation.

Secondly, as we select a small subsample, the overall coverage of every method decreases as compared to the full cohort. This, of course, is to be expected because a partial set of studies inevitably leaves out a significant amount of information. Interestingly, when we include more studies in our benchmark, the lead of BiGER and gBiGER becomes more noticeable, and the performance delta between methods becomes larger. In the real-world setting, it is often the case that hot topics in the literature result in large meta-analyses with many studies, and the advantage of BiGER is thus a practical one. Conversely, the scenario of a subsample size of 3 results in all methods clustering together, although BiGER is still at the top. This can be explained by the fact that this situation reaches the lower limit of meta-analysis: in fact, meta-analysis needs at least two studies by definition. Nonetheless, the BiGER family’s lead in this scenario shows its robustness in the extreme scenario with a minimally small subset of studies in literature.

### BiGER is Robust Against Study Selection

As part of the meta-analysis workflow, one of the main steps is to select relevant studies to be included for rank aggregation. While some studies’ inclusion can be obvious, tangentially related topics and experiments sometimes create ambiguity for researchers. The question, therefore, becomes whether researchers should err on the side of over-inclusion, or they need to focus on a relatively specific set of results. Luckily, BiGER’s posterior estimate for study variance 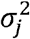 can shed light on genes’ deviation from their overall importance in study *j*, thus assessing the relevance of each study.

To quantifiably assess BiGER’s variance estimation performance and its usefulness in helping researchers select studies, we take a twofold approach for analyses. First, we simulated a situation where irrelevant studies are included by adding randomly generated studies to the default simulation procedure described above. Given the data-generating mechanism of our simulation, randomly generated studies should have minimal to no contribution to the estimation of truly important genes. Specifically, we included two scenarios: one with 3 irrelevant studies and one with 5 irrelevant studies. The latter represents a more extreme situation, for effectively one third of the studies in each cohort is random.

Inspecting the posterior estimates of variance of all studies in such cohorts, we found that both variants, BiGER and gBiGER, can successfully identify irrelevant studies (**Fig. 3b**). Specifically, we performed RA with both relevant and irrelevant studies and found the top 3 or 5 studies with the largest study variance based on posterior estimates. The procedure was repeated 25 times to aggregate all the results. Our results show that irrelevant studies (the left violin plot for each method) indeed have the largest variance within each cohort estimated by BiGER. In fact, the difference is quite obvious, considering the log scale of the y-axis. This suggests that a data-driven approach will work well for practitioners to identify anomalies in their meta-analysis. gBiGER, on the other hand, exhibits an interesting pattern of extremely large variance combined with a few studies whose variances are mixed with informative ones. This is likely due to the fact that the boundary approximation of BiGER induces a more constrained latent space, limiting the behavior of the variance. In this case, fixing the boundary of *ω*_*gj*_ as an approximation scheme proves to be even more advantageous because it can clearly delineate study’s relevance. Meanwhile gBiGER is still a useful tool in identifying most erroneously included studies. The variance estimation procedure thus provides a guideline for researchers to identify anomalies and conduct post-hoc analyses to identify the reasons.

While obtaining the posterior estimate of study variance is useful, it is also important to assess the robustness of BiGER if irrelevant studies were to be included anyways. Using the same simulation cohorts, we repeated the experiments by removing the irrelevant ones and computing the coverage as an accuracy metric. Strikingly, BiGER does not see a significant performance drop with irrelevant studies included, whereas MAIC sees a larger drop in performance (**Fig. 3c**). gBiGER sees a small difference in performance, but its coverage is overall on par with that of BiGER. This result is a good indication that BiGER is robust against noise in the study selection procedure. The same observation can be made for BiGER when using the Spearman correlation as a metric (**Fig, 3d**). MAIC’s performance seems to increase with irrelevant studies, which is interesting but somewhat counterintuitive and so hard to explain.

Besides completely irrelevant studies, another foreseeable scenario consists of studies with discordant data-generating mechanisms. Namely, we assumed *μ*_*g*_ ~ *N* (0,1) in our model, but in reality, many studies’ rankings are generated using latent processes specific to each study’s design and analysis pipeline (*e*.*g*. differential expression analysis, experimental validation, *etc*.). We thus further included a simulation scenario in which studies were relevant but violated the underlying distributional assumption of BiGER. Specifically, we tested the following distributions^6^ for generating *μ*_*g*_: 1) An exponential distribution with the rate parameter of 1, denotated as *Exp*(1), for a positive, right skewed ranking; 2) A folded *N*(0,1) distribution to simulate results from hypothesis testing; 3) A *t* distribution with 3 degrees of freedom with heavier tails. For the latter two settings in which the variance is not 1, we standardized the sample with its standard deviation. Using both coverage and Spearman’s correlation as metrics, we found that the BiGER family is still the most accurate among the methods tested (**Sup. Fig. 1**). This result shows that the BiGER family is robust against departures from assumptions on the latent variable model. Considering the fact that BiGER already performs well across a range of settings with studies of different quality, it is reasonable for researchers to include as many relevant studies as possible without sacrificing performance.

### BiGER Is Consistently Accurate in Real Benchmark Studies

Having validated BiGER’s posterior ranking performance and variance estimation using simulation, we next turn our focus to real datasets in the literature. A large-scale meta-analysis is a nontrivial task, for curating and interpreting a large cohort of studies requires the collaborative effort of domain experts. Luckily, there exists three large benchmark studies that have previously been proposed for assessing the accuracy of aggregating both top-ranked and top-unranked lists^5,6,16^. Following the original naming scheme, these three cohorts are named as the following: Macrophage, NSCLC, and SARS-COV2. The Macrophage cohort consists of 4 ranked studies and 2 top-unranked studies; the NSCLC cohort has five top-ranked studies; and the SARS-COV2 cohort has 11 top-ranked studies and 21 top-unranked studies. The characteristics of the three cohorts are summarized in more detail **Sup. Table 1**. We then performed RA with both variants of BiGER along with other methods for benchmark, and we used the provided truth set for computing the coverage statistic. Here, all unreported genes in each list are treated as bottom ties in accordance with previous recommendations and assumptions^6,16^. We included details of the preprocessing procedures and their rationales in the **Methods** section.

As shown in **Fig. 4**, we found that both BiGER and gBiGER have relatively good coverage across all settings in all three cohorts. Besides investigating coverage itself, we also focused on the difference between the top method and the remaining methods. We thus calculate the coverage of a given method as a percentage of the top method within a given setting and cohort, which are shown as colors in the plot. It is then obvious that BiGER performs especially well for the Macrophage and SARS-COV2 cohort, whereas the gBiGER is one of the top contenders for NSCLC. Despite BiGER’s lead in the Macrophage cohort, gBiGER still outperforms methods outside of the BiGER family regardless. No other method performs well across all three cohorts. The original BiG performs poorly in the Macrophage and SARS-COV2 cohorts, both of which contain top-unranked lists, highlighting the need for specifically modeling them. MAIC is a good contender for the NSCLC and SARS-COV2 cohorts, but it does not come close to the coverage of the BiGER family in the Macrophage cohort. Simple methods, such as those from the Borda’s collection, do not possess the prowess of producing competitive coverage under the default setting.

**Fig. 4.**
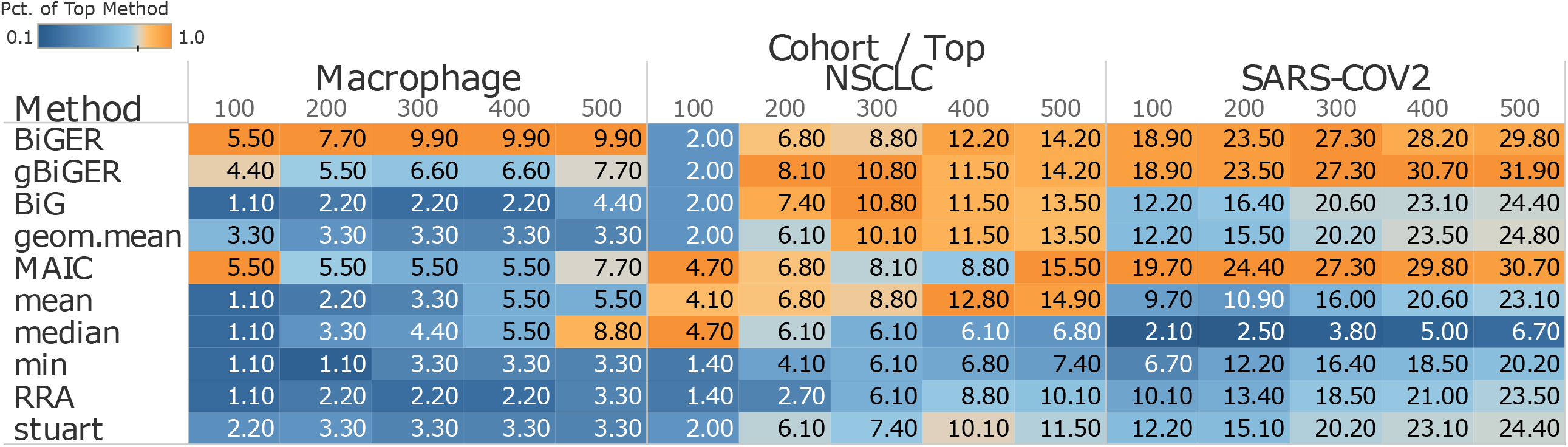
The performance of BiGER in three real benchmark datasets. A comprehensive evaluation of the BiGER family and the other methods with the Macrophage, NSCLS, and SARS-COV2 datasets. All numbers in the table represent coverage. The color in each cell is based on the percentage of the top performing method in the corresponding setting. Orange color indicates overall superior performance at 100%, whereas blue color means deficiency against the top method.

NSCLC is the only cohort without top-unranked lists, and interestingly, no method has a clear advantage here. Methods such as arithmetic mean and MAIC perform just as well as BiGER. The reason is twofold. First, the lack of top-unranked lists allows all methods, including BiG, to utilize as much information as available in the dataset. Second, this cohort was originally curated by Li et al.^5,6^ with the intention to model platform effect as well. Specifically, the datasets consist of sources from both bulk RNA-seq and Affymetrix microarrays. The benchmark dataset was further recompiled by Wang et al.^16^ to include a subset of the studies for which the platforms are not specifically given. The results here therefore are concordant with previous results in the literature but with the stated caveats. Nevertheless, the wide adoption of this dataset provides a good platform for validating BiGER and its benchmarking procedures (See **Sup. Fig. 2** for convergence properties showcased using the NSCLC cohort and **Methods** section for detailed benchmark settings).

In the Macrophage and SARS-COV2 cohorts, the performance gap between the leading methods and the rest are much wider. BiGER dominates across the board in the Macrophage cohort, with gBiGER lagging slightly behind. Given that the Macrophage is a real cohort with only 91 genes in the truth set, BiGER’s dominance in this dataset is an important validation for its performance: a large coverage difference in a relatively small cohort validates BiGER’s ability to identify a few more truly significant genes, which are useful for practitioners who wish to use RA results for further research. For the SARS-COV2 cohort, BiGER, gBiGER, and MAIC performed the best. In fact, these three methods are the only ones that specifically support unranked studies. Given that this cohort is largest of the three with 32 individual studies and a large portion of them being unranked (11 ranked and 21 unranked), this benchmark provides good evidence for the advantages of specifically modeling unranked lists.

Beyond coverage as an overall metric for assessing BiGER’s accuracy, its study-specific variance estimation is also an important tool for understanding individual studies and validating the meta-analysis process. The large-scale nature of the SARS-COV2 cohort not only reflects the amount of literature recently published on the topic, but it is also naturally suited for this task. By comparing the posterior estimates of study variance of all 32 studies (**Sup. Fig. 3**), we found that studies with the smallest variance are large, ranked studies, whereas unranked studies have slightly larger variance. This trend makes a logical explanation for the information carried in each study: top-ranked studies have precise information about the ranking positions, and at the same time, those with higher true coverage are more concordant with the truth set. Further, we found that there is no study with abnormally large variance, suggesting that the benchmark dataset consists of relevant studies with a few that are particularly important. This analysis again showcases the versatility of BiGER in the real setting, allowing researchers to perform RA on all reasonably selected studies and conduct *post-hoc* validation using estimated study-specific variance.

### Efficient Inference via Variational Inference

One of the core contributions of the BiGER family includes a VI-based algorithm to improve a Gibbs sampler’s efficiency. The original Bayesian hierarchical method from BiG and its corresponding implementation suffer from poor efficiency. In fact, it is necessary to truncate gene lists by excluding genes that are not top-ranked in any study to speed up computation in their original benchmark^6^. BiGER, on the other hand, makes vast efficiency improvements by implementing VI from the ground up in RCPP^30^ with careful code optimization. To show the performance gain, we first conducted a small-scale efficiency benchmark by comparing the runtime of BiG to that of the BiGER family. We found that gBiGER is two orders of magnitude faster than BiG, whereas BiGER further improves upon gBiGER by more than one hundred times (**Fig. 5a**). Importantly, the default setting consists of only 10 studies with 1000 genes, which is a rather moderate meta-analysis in genomics. Our benchmarking process has revealed that BiG is increasingly slow with larger gene lists, and we thus restricted the scope of our first benchmark.

**Fig. 5.**
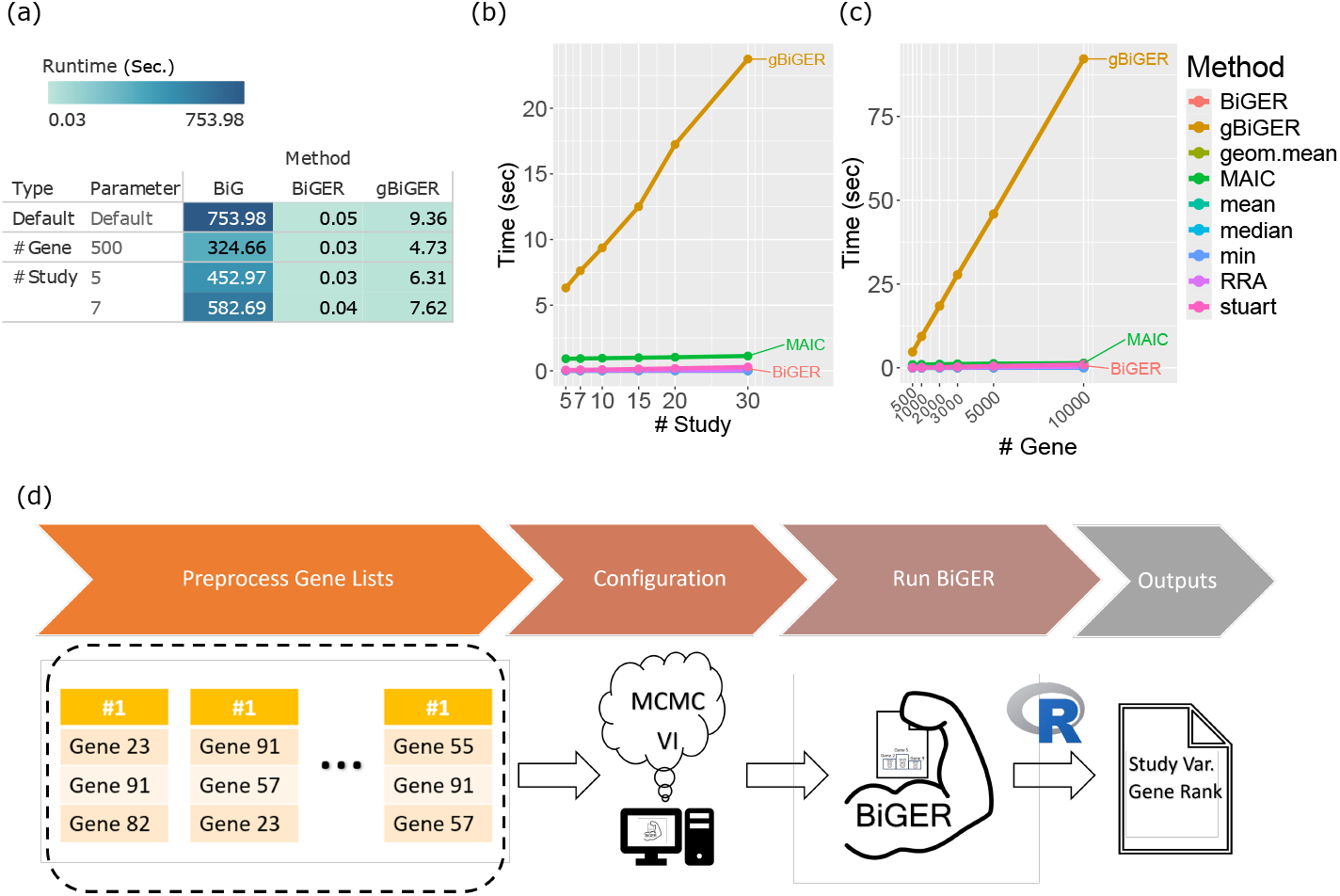
Efficiency and usability of BiGER via a software pipeline. (a) A comparison of runtime between BiG and the BiGER family. (b)-(c) Runtime benchmarks of the BiGER family against other methods, except for BiG. The number of studies in (b) and the number of genes in (c) are compared. All runtimes are measured in wall-clock seconds. (d) A practical pipeline for executing the BiGER’s software from collecting studies to obtaining posterior ranking results. The R logo is licensed under CC-BY-SA 4.0 from the R Foundation.

Moving beyond BiG, we proceeded to test the BiGER family against other methods with a variety of settings. We first varied the number of studies from 5 to 30, and we found that gBiGER’s runtime increased linearly (**Fig. 5b**). While this is to be expected, other methods can be run in almost-trivial time. Encouragingly, we saw that BiGER is faster than MAIC, and its runtime almost approaches that of the Borda’s collection. Although simple statistics-based methods in the Borda’s collection are technically faster, the difference on the scale of a fraction of a second is negligible. On the contrary, gBiGER’s relative slow runtime here is explained by the fact that a Gibbs sampler needs a reasonable number of iterations to converge, whereas BiGER’s MFVI can converge in far fewer iterations.

We then moved onto benchmarking the effect of genes by increasing the number of total genes considered. Again, the same pattern can be observed (**Fig. 5c**). Due to scaling, methods including BiGER and MAIC all can be run efficiently with BiGER posing a slight advantage. The efficiency benchmarks in **Fig 5a-c** show that gBiGER offers vast efficiency improvements over BiG, whereas BiGER can be considered trivially fast. Interestingly, even under the setting with 10,000 genes, gBiGER takes less than 2 minutes, which is reasonable. For most researchers, BiGER, which combines efficiency with good accuracy, is our recommendation. Such efficiency can be crucial when working with massive datasets with the possibilities of extending BiGER to other fields in the future. For those who need the theoretical guarantee of a Gibbs sampler and are working with reasonably sized datasets, gBiGER is a good choice.

### A software package for rank aggregation

For practitioners in the field, both BiGER and gBiGER proposed in this work offer model-based RA in a Bayesian meta-analytic framework. Practically, we foresee two important workflows, both of which we support natively in an easy-to-use R package with C++ backend **(Fig. 1a**; **Methods** section). In a typical meta-analysis, researchers compile gene lists from scientific literature and then perform RA as the second step. In this workflow, BiGER allows researchers to directly input gene lists as a list object in R, and we offer a convenient preprocessing function to appropriately process them in accordance with BiGER’s model specification. For example, users can specify which gene lists are top-ranked, top-unranked, or mixed. Further, there is the option to treat unreported genes as either bottom ties or NAs, giving users the choice based on their needs and their understanding of the studies at hand. A diagram of an example workflow is shown in **Fig. 5d**. For the second workflow, some researchers and practitioners need to perform RA based upon a series of experiments or assays of their own. This data analysis pipeline typically includes differential analysis to obtain important genes for a given condition, which are then used for RA. One of the most common differential analysis tools in R is Seurat, and it does not output a gene list by default. To address this, we implemented a preprocessing step to extract gene lists from Seurat outputs, which are subsequently compatible with BiGER.

As a general Bayesian RA framework, it is important for users to easily perform diagnostics and understand the MCMC run. For all gBiGER runs, we output posterior estimates for both ***μ*** and ***σ***^2^. Users have the option to save the entire MCMC chains with or without the burn-in period such that they can easily produce trace plots or analyze data in popular downstream pipelines, such as that of the *posterior* package^31^. Running multiple chains is also supported natively without needing to call BiGER repeatedly. For BiGER, we output both and variational estimates for ***μ*** and ***σ***^2^ the convergence criterion *δ*, whose definition is included in the **Methods** section. All outputs are presented in a list objects, which are easily compatible for downstream analysis and diagnostics. It is key to note that all variational distributions in BiGER are known in closed form for all parameters, allowing for uncertainty quantification through their variances easily. Thus, it is possible to perform interval estimation and posterior inference.

## DISCUSSION

In this work, we propose a family of Bayesian RA methods with support for extended ranking schemes. Combining a rigorous statistical framework with VI, BiGER and its variant gBiGER achieve not only good accuracy but also competitive efficiency. The proliferation of high throughput sequencing technologies such as scRNA-seq and the ease for researchers to compile relevant studies present a need for integration methods. More pressingly than ever, it is necessary to gauge the confidence in published literature and find a consensus upon which downstream treatment and policy-making can rely. BiGER therefore fills this role by offering a robust algorithm with interpretable study-specific variance as a practical solution for researchers.

BiGER, as compared to other methods in the field, poses a few key advantages. First, it offers the most flexibility in the ranking scheme without making *ad-hoc* assumptions. To our knowledge, BiGER is the first method to support mixed studies with both top-ranked and top-unranked genes. From the data model, it is trivial to extend the existing framework to allow arbitrary ranking schemes, such as inclusion of multiple groups of ties. Such flexibility is not present in MAIC. While methods in the Borda’s collection can in theory support any input matrices, *ad-hoc* preprocessing is oftentimes necessary in the presence of ties. Specifically, the method used to assign ranks to tied items (*e*.*g*. the mid-rank method and top-rank method among others^13^) is up to each user’s discretion. BiGER, on the other hand, treats all input ranking as is, which is its key advantage to achieving reliable and repeatable inference. Second, BiGER, along with BiG, are the only methods tested to report study-specific variance. As shown in our numeric results, study-specific variance carries important information on the quality and nature of the input studies. BiGER thus offers the benefit of interpretability beyond a simple aggregated ranking.

Beyond what we tested, BiGER also serves as a foundation for future models in meta-analysis. New sequencing technologies will inevitably arise. Along with them comes the need for more flexible designs specific to each meta-analysis. The possibilities of extension, we argue, lie upon the specific circumstance rather than a one-size-fits-all solution. For example, one meta-analysis may need to account for platform effects, whereas another may involve medical conditions from different geographic regions. We therefore note that BiGER’s basis on a Bayesian linear model allows an easy addition to incorporate additional random effects or covariates, where a VI algorithm can be similarly derived. For most users right now, BiGER already offers the most flexible solution to date; for others requiring more complex modeling with additional parameters, BiGER provides a good starting point upon which to build a customized model.

From a broader perspective, BiGER is situated in the RA literature with many applications in both statistics and beyond. Despite the focus on genomics in our study, it is paramount to realize that many of the concepts here can be translated into general applications: each study corresponds to an input list and each gene corresponds to an item of interest. The assumptions on input lists can be easily relaxed for accommodating any tie patterns in the general sense. We posit that the Bayesian framework in BiGER is indeed a good fit for aggregating data in the form of ordinal lists with similar assumptions. In the RA literature, there is a common thinking that trivial methods such as arithmetic or geometric mean are sufficient. BiGER’s usability and extensibility, however, present a good case for its wider adoption in the future. It is our hope that this work will serve as a solid foundation upon which future works will flourish.

## METHODS

### Genes and Ranking Schemes

Given the complexity of notations introduced in this paper, a full list of notations and their associated definitions are included in **Supplementary Note 1** for reference. Before further expanding the statistical details of the BiGER model, we first need to rigorously define all possible ranking schemes within our framework by considering all types of genes based on their available ordinal information. Given the full set of genes 𝒢 in a given meta-analysis, we categorize them into one of the four categories: 1) top-ranked genes, 2) top-unranked genes, 3) bottom-tied genes, and 4) missing genes (NA’s). As previously discussed in the **Results** section, both top-ranked and top-unranked genes are considered observed and thus reported by a given study. Bottom ties, on the other hand, may be studied but unreported, and their status needs to be deduced based on whether each study formally investigated these genes. Missing genes are as truly missing in the sense of missing data, possibly due to issues such as sequencing quality control.

By definition, when present in an input list, top-ranked genes are considered the most important, followed by top-unranked genes and bottom-tied genes. For missing genes, however, no information is available regarding their relative importance. Considering all four types of genes, we theoretically have 2^4^ or 16 total combinations for possible ranking schemes. Their compositions are detailed in **Sup. Table 2**. However, not all combinations can be modeled. First, all eight cases with top-ranked genes are valid within our framework. This is a straightforward extension of the ranking framework proposed by Li et al.^6^ by including four cases with top-unranked genes. Second, in the absence of top-ranked genes, only two cases with the presence of bottom ties are nontrivial in practice; the other six situations have only either top-unranked or bottom ties and thus have no ordinal information available at all. In other words, for top-unranked studies, we require the presence of bottom ties. The reasoning is rather simple: our likelihood is formed by modeling the ordering of genes or groups of genes. If you only have a single group of ties, either bottom ties or top-unranked, we have no information on their relative importance. For example, a single group of top-unranked genes has the likelihood of

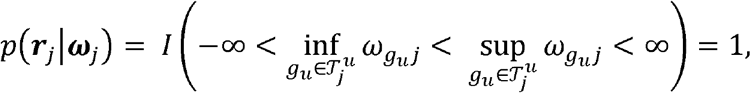

which carries no practical insight.

Having considered all valid cases of ranking schemes, we can consolidate 10 of these schemes into three general types of lists. Namely, we define these lists with the following necessary conditions:

1. Mixed list: 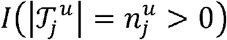 and 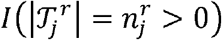

3. Top-unranked list: 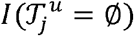, and 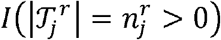.

2. Top-ranked list: 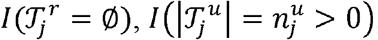 and 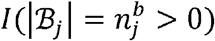

For mixed lists and top-ranked lists, it is possible and in fact likely to have bottom ties, but we do our model. not impose it as a condition. We thus use these indicator functions to identify each type of list in our model As a reiteration, we use the notation 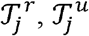, and ℬ_*j*_ to represent the set of top-ranked genes, top-unranked genes, and bottom ties. Their cardinalities are 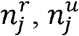, and 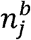 respectively. In a given study *j*, we define the number of relevant, non-missing genes as 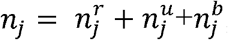, and all such relevant genes have likelihood contributions in our model. Missing genes, on the other hand, provide no information and thus naturally disappear in our likelihood for an all-available case analysis.

### Full BiGER Model

As shown in the **Results** section, the full probability model of BiGER, *p* (***r, Θ***), can be written in the form of Eq. (1), where ***r*** are observed ranks and is **Θ** the collection of all parameters. One of the most intricate parts of this model is *p* (***r***, | ω), which is nontrivial despite the conceptual simplicity of the order being correct.

For the simplest case, let us first consider top-ranked lists, which are conventionally the easiest case. In the presence of bottom ties, the ordering of latent weights is

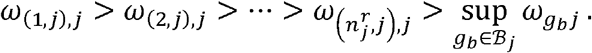

We consider bottom ties as a single group with pooled boundaries in which only the largest element will affect the ordering. However, complications occur when there are no bottom ties because ℬ_*j*_ = Ø makes the definition of the supremum unclear. To address this issue while keeping the definition concise, we define

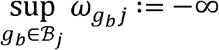

when ℬ *j =* Ø, in the absence of bottom ties.

Next, we consider top-unranked studies in which there are no top-ranked genes. The only requirement in this case is that all top-unranked genes have weights that are higher than bottom ties, which definitely exist as previously discussed. Like bottom ties, top-unranked genes are conceptual ties, and in this case, only the smallest weight will affect the ordering. The condition can therefore be written as

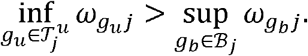

For mixed lists, it can be conceptualized as a combination of top-ranked and top-topunranked lists, where bottom ties are again optional in this caseas Given the flexibility of 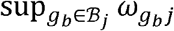 as defined previously, we can simply combine the conditions of top-ranked and top-unranked lists to form the condition for mixed lists:

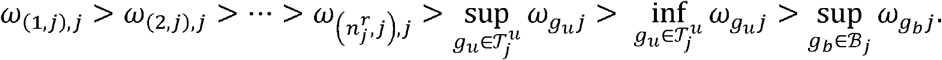

Having defined the conditions for all three types of lists, we have all the necessary ingredients for *p* (***r***│***ω***). Putting everything together, the likelihood can be written as:

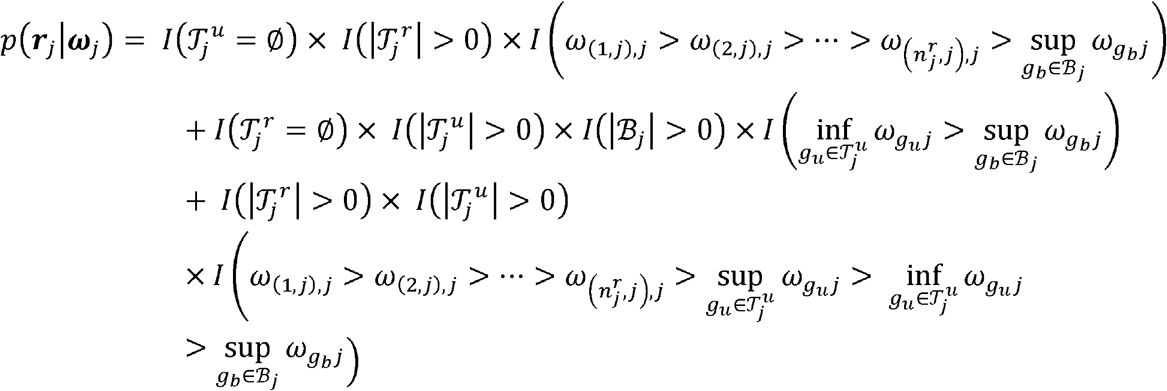

Note that this rigorous expression is consistent with the key intuition that the order of latent weights is correct. *p* (***r***_*j*_ │ ***ω***_*j*_). consists of a series of indicator functions, where each study falls under three mutually exclusive conditions.

The full form of *p* (***r***_*j*_ │ ***ω***_*j*_) is a bit cumbersome with the explicit consideration of three different types of lists. In practice, we can simplify it (with a slight abuse of notations) by considering top-ranked and top-unranked lists as special cases of mixed lists. Then, we can write *p* (***r***_*j*_ │ ***ω***_*j*_)as

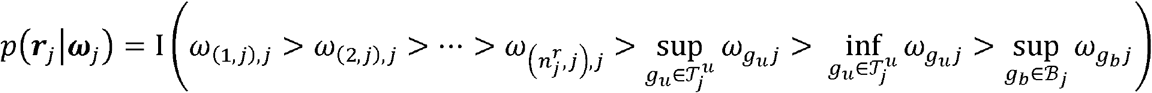

with the caveat that only the relevant terms appear for each list. Using this simplified expression, we can write down the full probability model:

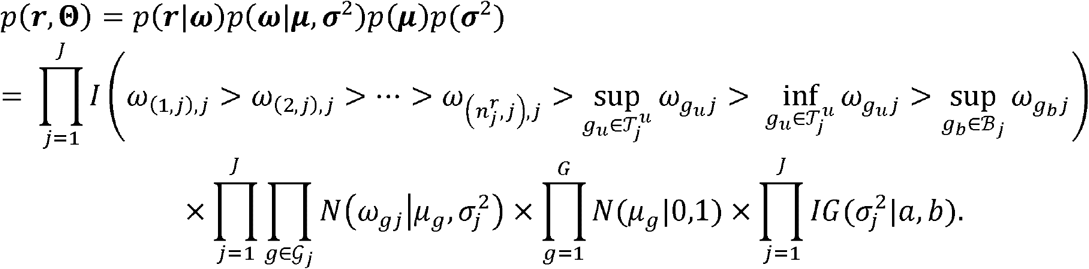

With the posterior distribution proportional to the above full model, we proceed to derive a Gibbs sampler for posterior inference.

### Gibbs Sampling for gBiGER

The full conditional distributions for *μ*_*g*_ and 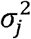 are relatively straightforward to derive since they do not involve the indicator function. They are

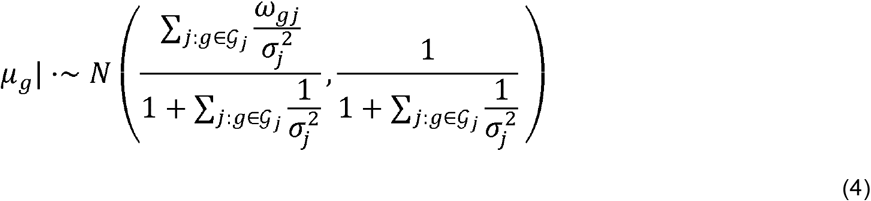

and

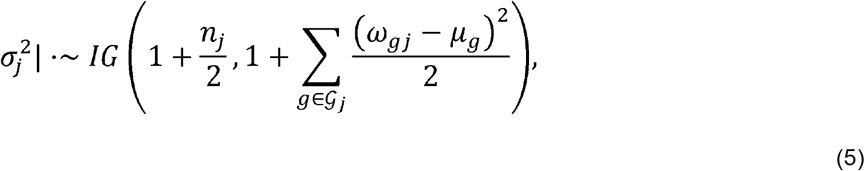

where *n*_*j*_ is the number of relevant, non-missing genes in each study as previously defined. A note of notation and implementation here is that we sum over all the genes present in each study *j*, accounting for truly missing genes by assumption.

Using the full model with the simplified indicator function, we can derive the full to be conditional distribution of *ω*_*gj*_ to be

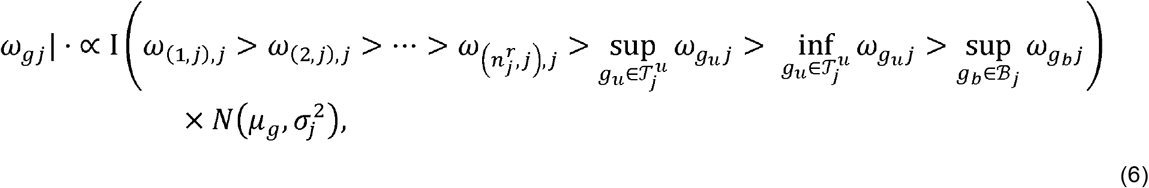

which is a truncated normal (TN) distribution. However, the truncation bounds are unclear using this form because bounds depend on the specific ranking of *ω*_*gj*_ and its ranking nature. We thus derived a more specific algorithm for sampling *ω*_*gj*_. Given that all TN distributions are based on 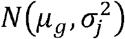, we from now on focus on the truncation bounds. Intuitively, the procedure is to pick *ω*_*gj*_ and its neighboring items from the indicator function, which forms a proper density for the TN distribution. To more rigorously consider the bounds, we treat different types of genes separately.

First, we consider top-ranked genes whose latent weights can in general be sampled with the lower and upper bounds of 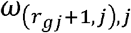 and 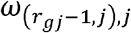. There are a few special cases we need to consider. The gene with *r*_*gj*_ = 1 in a given study has the upper bound of ∞. On the other hand, the gene with 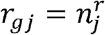 has a lower bound based on the following conditions: if 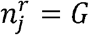, which means that the study consists of a full top-ranked list, then the lower bound is set to − ∞; if there exists top-unranked genes, the lower bound is 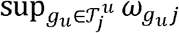; otherwise, there are only bottom ties, which implies a lower bound of 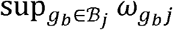.

Secondly, we consider top-unranked genes. Here, all the genes can be sampled in bulk because they share the boundaries. When there are top-ranked genes, the upper bound is 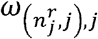, which is the latent weight of the last-ranked gene in the study; otherwise, the study consists of a top-unranked list with an upper bound of ∞. Since all unranked genes have larger latent weights than those of bottom ties, the lower bound is 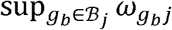

Thirdly, the boundaries for bottom ties are similar, and they can again be sampled in bulk. When there are the top-unranked items, the upper bound is 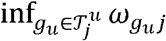; without top-unranked genes, the upper bound is 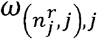 The lower bound of bottom ties is always − ∞. Note that the upper bound for bottom ties can never be ∞ by assumption because this will imply that there are lower bound of bottom ties is always by assumption because this will imply that there are no top-ranked and top-unranked genes, consisting one of the invalid cases shown in **Sup. Table 2** for reasons previously discussed.

The implementation of a Gibbs sampler needs some care, especially regarding the sampling step for *ω*_*gj*_. Namely, at each iteration, the order of which top-ranked genes’ latent weights are sampled is randomized. This is to prevent drifting: given that the genes are ordered, a sequential approach for sampling from the highest ranked gene onward will skew the weights of all genes based on the first-sampled value. A randomization procedure with sufficient number of iterations will solve this issue.

The overall Gibbs sampler can be summarized with the following algorithm:

1. Initialize ***μ, σ***^2^, and ***ω***
2. For *j* in in 1,…,*G*, draw *μ*_*g*_|. from the normal distribution given in (4).
3. For *g* in 1,…,*J*, draw 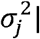 from the inverse gamma distribution given in (5).
4. For *j* in 1,…,*J*, sample *ω*_*j*_ using the following steps (skip if specific gene types are not present in the study):
  a. Randomize the sampling order of top-ranked genes. For *g* in 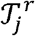, draw ***ω***_*gj*_ from the truncated normal distribution given in (6) with the appropriate boundaries according to the order.
  b. Draw 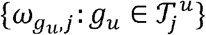 from the truncated normal distribution with the appropriate boundaries.
  c. Draw 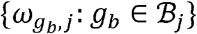 from the truncated normal distribution with the appropriate boundaries.

### Variational Inference for BiGER

In the full BiGER model, we derive a VI algorithm for efficient parameter estimation. As previously shown in Eq. (2), the goal of the VI is to derive a family to independent variational distributions *q* (·) to approximate each parameter. We employ the Coordinate Ascent Variational Inference (CAVI)^29^ algorithm to find *q* (·), which iteratively maximizes the ELBO via Eq. (3). While a typical conditionally conjugate model does not pose a challenge for CAVI, the indicator function in BiGER’s likelihood is nontrivial for optimization. Namely, the boundaries of the TN distributions for *ω*_*gj*_ can change in each iteration. This parameterization works well for gBiGER because the computation of full conditionals does not require expectations, and thus we can derive TN distributions in closed form. However, this is not the case for CAVI. More specifically, were we to derive BiGER directly, we need to find the expectation

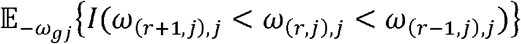

where, without loss of generality and for the sake of demonstration, we assume 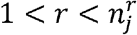 here. We have thus developed an approximation scheme to address this issue. The rationale and details of this derivation are further detailed in **Supplementary Note 1**.

The approximation scheme is rather simple: for *ω*_*gj*_, we define a concordant and known boundary *a priori* to avoid the need of depending on other parameters. This is achieved via a 1-1 transformation of the observed ranks. Given that the scale of the latent space does not matter, there are many choices of valid transformations. Because of the fact that *μ*_*g*_ follows a normal fact that the Normal quantile function maps distribution with support on the real line, it naturally makes sense to employ Normal quantile this approximation. We start by noting the obvious fact that the Normal quantile function maps for [0,1] to the real line. We thus need to partition according to the correct ranking schemes to each study to obtain the quantiles. The exact procedure is as follows:

1. We partition the unit interval [0,1] into *n*_*j*_ equally-spaced sub-intervals using *n*_*j*_ − 1 points (excluding the end points).
2. The top-ranked genes fall into the top 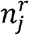 intervals in order (*e*.*g*. the first top-ranked gene has the upper bound of 1 and lower bound of 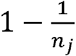, and so on and so forth).
3. For top-unranked genes, the intervals of the 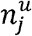 unranked genes are pooled with an interval length of 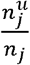 and correctly position in relation to top-ranked genes and bottom ties. The procedure is the same for bottom ties.
4. All missing genes in each study are not assigned boundaries.

For gene types that do not exist in some studies, the intervals are shifted accordingly analogous to the scheme in **Fig. 1b**. It is key to point out that even though this scheme seems similar to that of the gBiGER procedure, the bounds remain fixed throughout the algorithm.

Using this approximation, the likelihood becomes *p* (***y***_*gj*_ | *ω*_*gj*_), where *y*_*gj*_ consist of 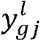 and 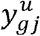, which are the lower and upper bound for each gene. *y* is treated as data just like the observed ranks ***r*** in the BiGER model. Therefore, the expectation involved in CAVI is

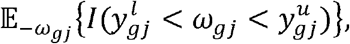

which no longer poses a concern because *ω*_*gj*_ is the only parameter (more details in **Supplementary Note 1**). After some derivations, we arrive at the variational distributions we show below. For convenience, all expectations shown are taken with respect to the enclosed parameters. The variational distributions are

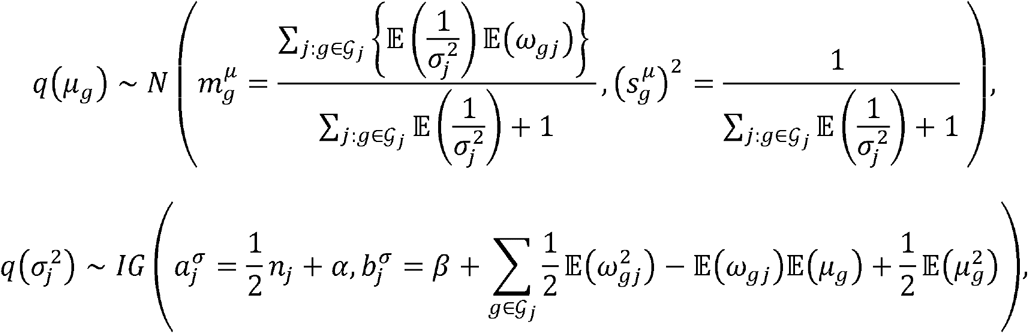

and

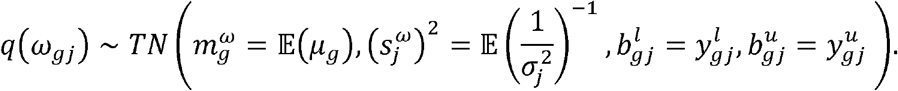

In each iteration, we update the parameters of the variational distributions directly using the formulations here. All expectations here can be computed in closed-form, and they are included in **Supplementary Note 1**. There is no need to randomize the order of top-ranked genes while updating 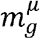 because of fixed bounds. VI-approximated posterior ranking is computed by ranking *m*^*µ*^ in the last iteration.

### Simulation model overview

To comprehensively validate the BiGER family of methods with known ground truth, we devised a simulation model based on the latent variable assumption. In general, the simulation is to be done in two stages. First, we need to simulate the meta-analysis specification and study composition. Given that real studies consist of different types of lists with unique features, there is a need to realistically simulate these lists with settings such as proportion of ranked or unranked items, length of the lists, etc. Second, once the specification of a given meta-analysis is simulated, we need to generate the ranking both globally as the ground truth and locally for each list. Here we thus describe each stage in detail.

In the first stage of the simulation, we specify the total number of genes *G* and the total number of studies *J*.*J* is usually known in the real setting when a literature search has concluded, whereas the upper bound of is determined by these studies. To accurately simulate the composition of a cohort with a mix of mixed, top-unranked, and top-ranked studies, we sample the type of list with a categorical distribution with the corresponding event probabilities of 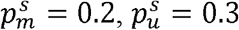, and 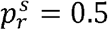. For a given study, the next step is to determine the number of each type of genes present. Recall that there are four types of genes: top-ranked, top-unranked, bottom ties, and missing genes. The distribution of these genes in each study follows *Multi*(1,*G*,***p***^*g*^), where ***p***^*g*^ ~ *Dirichlet* (***α***) In other words, we sample the number of each type of gene using a multinomial distribution, whose event probability vector follows a Dirichlet distribution with the concentration vector ***α*** = {*α*_*r*_, *α*_*u*_, *α*_*b*_, *α*_*NA*_} that needs to be specified. The default simulation setting ***α*** of was set to{*α*_*r*_= 3,*α*_*u*_ = 2,*α*_*b*_ = 4,*α*_*NA*_ =1}. For studies without mixed or top-unranked lists, the corresponding terms in ***α*** are set to 0. Given that the missing genes are assumed to be randomly missing, they are sampled randomly in each study. The missingness parameter in the simulation cohort (**Fig. 2**) corresponds to the specification of varying *α*_*NA*_ ∈{0,2,3,4} with all other parameters in ***α*** held at the default value. The expected percentage of expected missing genes from the resulting multinomial distribution is simply 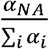 by the law of total expectation. An increasing value of *α*_*NA*_ increases the expected number of missing genes in each study, and vice versa.

Once all the study specifications are generated, we move onto the second stage by simulating the actual ranks in each study. We follow the general notations of BiGER with a latent variable model. The latent weight *ω*_*gi*_ is generated with 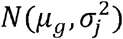, where *μ*_*g*_ ~ *N* (0,1) and 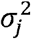 is generated via *ρ*_*j*_. We define *ρ*_*j*_ as the correlation between *μ*_*g*_ and *ω*_*gj*_ in a given study, and it turns out that 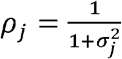 with the interpretation of *ρ* _*j*_ → 1 as 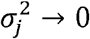 and *ρ*_*j*_ → 0 as 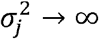. Intuitively, *ρ*_*j*_ controls the study quality within a cohort of studies. In our simulation (**Fig. 2**), *ρ*_*j*_ is either generated from *U*(*a*_*ρ*_, *b*_*ρ*_) or set directly. Once latent weights are generated according to this procedure, each study is ranked according to the weights.

As a note on technicality, all top-unranked and bottom ties are generated using the same assumptions of the latent variable model. Since *ω*_*gj*_ is inherently continuous from a normal distribution, we determine the membership of the 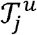 and ℬ_*j*_ by position using fully ranked *ω*_*gj*_. Specifically, for a mixed list, the first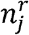 genes’ ranks are left as is for 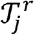 as top-ranked items; the items ranked 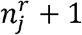 to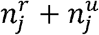 are logically denoted as top-unranked in 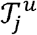; the rest of the ranked genes are bottom ties.

### BiGER Family Settings and Defaults

Both the Gibbs samplers and VI algorithm used in the BiGER family require some user-specified settings. In both our benchmarks and the software implementation, we use a set of standard parameters as defaults. Across all variants of BiGER, the Inverse Gamma prior for 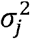 needs to be specified, and we set *a* = *b* =1 for a reasonably vague prior that allows the possibility of both large and small study variance (**Sup. Fig. 4a**). We performed a parameter tuning exercise using a range of IG parameters as previously suggested in the literature^6,32,33^ and found that there was no significant difference in coverage and spearman’s correlation (**Sup. Fig. 4b-c**).

The initial step of Gibbs update and CAVI both require initializations for ***μ, σ***^2^, and ***ω*** as
part of the algorithm. For simplicity, we sample ***μ*** ~ *N*(0,1) and set ***σ***^2^ = **1**, assuming no prior knowledge on gene ranking and study reliability. For ***ω***, the defaults depend on the algorithm of choice. In BiGER, boundaries for ***ω*** are fixed as previously described. Thus, we sampled each *ω*_*gj*_ from 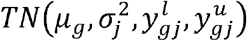 where initial values for *μ*_*g*_ and 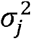 are used. For gBiGER, the weights for each study were sampled from *N*(0,1) and ordered so that they are concordant with the observed rankings. All these settings are also the default as implemented in the BiGER package, and they ensure that the initializations are vague without added bias.

gBiGER uses Gibbs sampling. By default, we perform 5000 iterations of sampling with the first 2500 steps discarded as the burn-in period. This achieves good convergence properties (**Sup. Fig. 2**) along with sufficient efficiency for practical use. The CAVI algorithm in BiGER requires far fewer iterations in general. We monitor the convergence by the absolute change in ***μ*** by the criterion of ‖ ***μ*** _*t*_ − ***μ***_*t*−1_‖ < *δ*, where we set *δ =* 0.0001 by default and subscript *t* denotes the number of iterations. We do not monitor the ELBO because the computation cost will be prohibitive. The defaults here result in fast convergence with good performance as seen in the benchmarks in the paper.

### Benchmark Implementation Details

The BiGER family of methods are implemented in-house with details in the software implementation section to follow. The BiG implementation was based on the Gibbs sampler with diffuse Inverse-Gamma prior as recommended in the original paper. For computation, we used an adapted version of the R codes shared by its authors (https://github.com/xuelilyli/BiG). The sampling procedures for the platform effect were removed for simplicity and efficiency, as platform effect is not considered in this paper. All defaults settings for BiG are the same as those for BiGER. For MAIC, we used the CLI provided by the authors (https://github.com/baillielab/maic) with default arguments. All other methods are part of the “RobustRankAggreg” R package, and we used the default settings for computation using gene lists as inputs.

BiGER is the only method tested that supports mixed lists, whereas all other methods require some preprocessing to handle mixed and sometimes top-unranked lists. The preprocessing is the most straightforward for MAIC, which supports both top-ranked and top-unranked lists. Mixed lists were converted to top-ranked lists by preserving only the top-ranked items. The lists were formatted in the suggested format in MAIC documentation. For BiG and methods in “RobustRankAggreg” package, all mixed lists were converted to top-ranked lists by treating top-unranked genes as bottom ties, whereas unranked lists were not included for computation due to its design limitations. BiG is the only method besides the BiGER family that distinguishes between bottom ties and truly missing genes. In real datasets where no such distinction is clearly provided, we treated all missing genes as bottom tie.

The efficiency benchmark of BiGER was based on the R version of BiGER with an RCPP backend. All R codes are timed with the “Sys.time” function and run in Windows 11. MAIC’s CLI is run as is and timed with Linux’s “/usr/bin/time” program under Ubuntu 20.04 LTS with Windows Subsystem for Linux. All computations were done on an AMD Ryzen 9 5900HX CPU with 32GB of DDR4 RAM.

### Statistics and Definitions

For all rankings in this paper, we assume a smaller rank to have superior importance, and ranks start at 1. For the BiGER family, we do not use any tie-breaking procedures given the models inherent capabilities in modeling top-unranked genes and bottom ties. In the evaluation of gene lists, coverage is defined as percentage of truly significant genes from the truth set that are covered in the top *K*_*TOP*_ genes after RA. For real datasets, the given truth sets are used; for simulation, the truth set is defined as the top 10% of the genes in a given simulation setting. By default, we set *K*_*TOP*_ = 100 but under a few benchmark settings, we varied *K* to include the set {100,200,300,400,500}, for a comprehensive understanding of coverage. Spearman’s correlation is implemented in R using the “cor” function with the “method= ‘spearman’” option.

In all trace plots, disperse starting points for ***μ*** and large initial variance ***σ***^2^ are selected, and all iterations including the burn-in iterations are plotted. All boxplots have boxes that extend from 1^st^ to the 3^rd^ quartiles with a line representing the median. Whiskers extend to the largest and smallest values within 1.5 times the interquartile range. Outliers are marked as dots.

## DATA AVAILABILITY

The public benchmark datasets are available at from Wang at al.^2^ (https://github.com/baillielab/comparison_of_RA_methods). The simulation datasets are generated using the BiGER software implementation in R (https://github.com/kevin931/BiGER).

## CODE AVAILABILITY

Both BiGER and gBiGER are implemented officially in R with extensive use of RCPP to provide efficient sampling and VI approximations. The R package offers a complete pipeline from preprocessing to gene lists, inference via BiGER, and downstream tasks. The R package for BiGER is available at https://github.com/kevin931/BiGER. The repository offers detailed tutorials on the package’s usage with examples and implementation details. Preprocessing pipelines are also included.

## AUTHOR CONTRIBUTION STATEMENT

KW performed simulations and real data analyses and developed the software. KW and XW developed the Bayesian methods and algorithms. XW conceived the main idea, designed and supervised the overall study. YY, YX, GX and JL provided feedback on statistical analyses and methodological development. All authors read and approved the manuscript.

## COMPETING INTERESTS STATEMENT

All authors declare no competing interest in this work.

## FIGURE AND TABLE LEGENDS

**Sup. Fig. 1.**
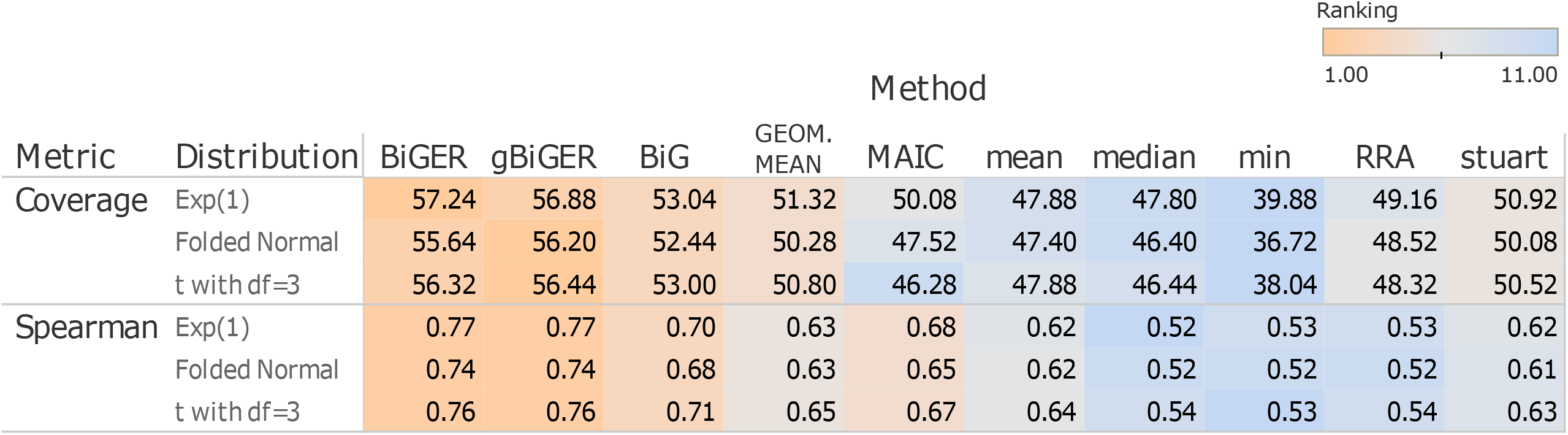
Robustness of the BiGER family against distributional assumptions. The coverage and Spearman’s correlation of RA methods with three different data-generating mechanisms are compared. Values represent the average coverage and Spearman’s correlation across all replicates for each setting, whereas colors indicate the ranking within each setting. Orange color indicates superior performance, and blue color represents inferior performance.

**Sup. Fig. 2.**
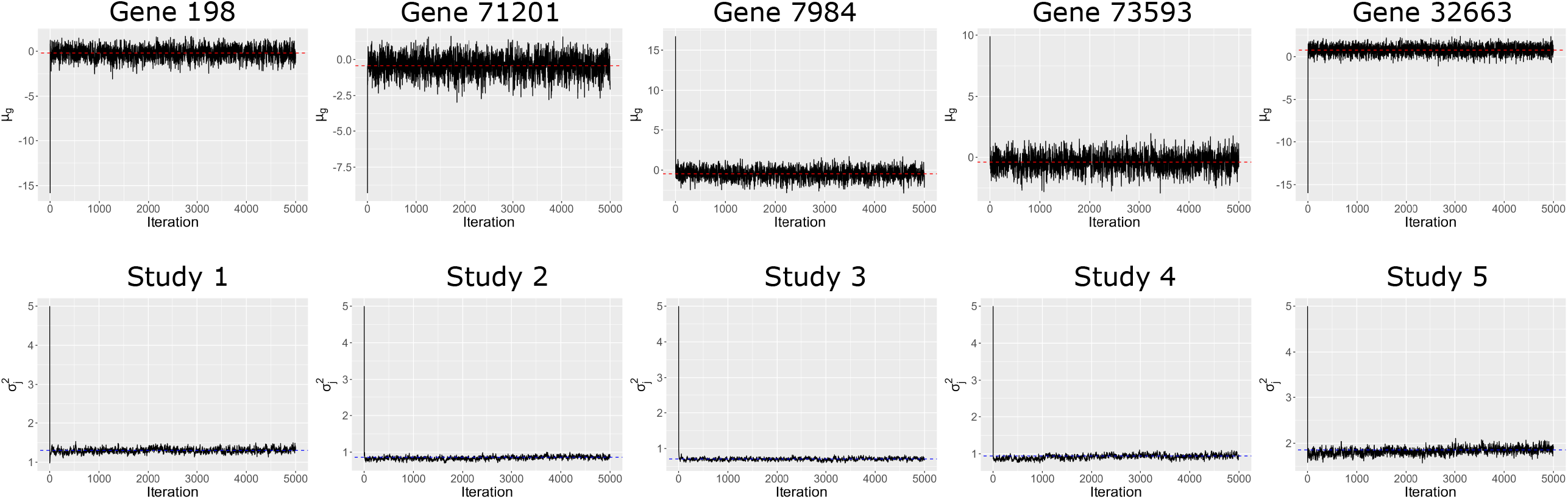
Convergence diagnostics of BiGER. The trace plot of *μ*_*g*_ for the 5 genes and 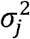 of all 5 studies of the NSCLC cohort are shown. The red dashed line indicates the posterior mean of *μ*_*g*_. The blue dashed line indicates posterior median of 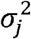. All iterations shown include the burn-in period.

**Sup. Fig. 3.**
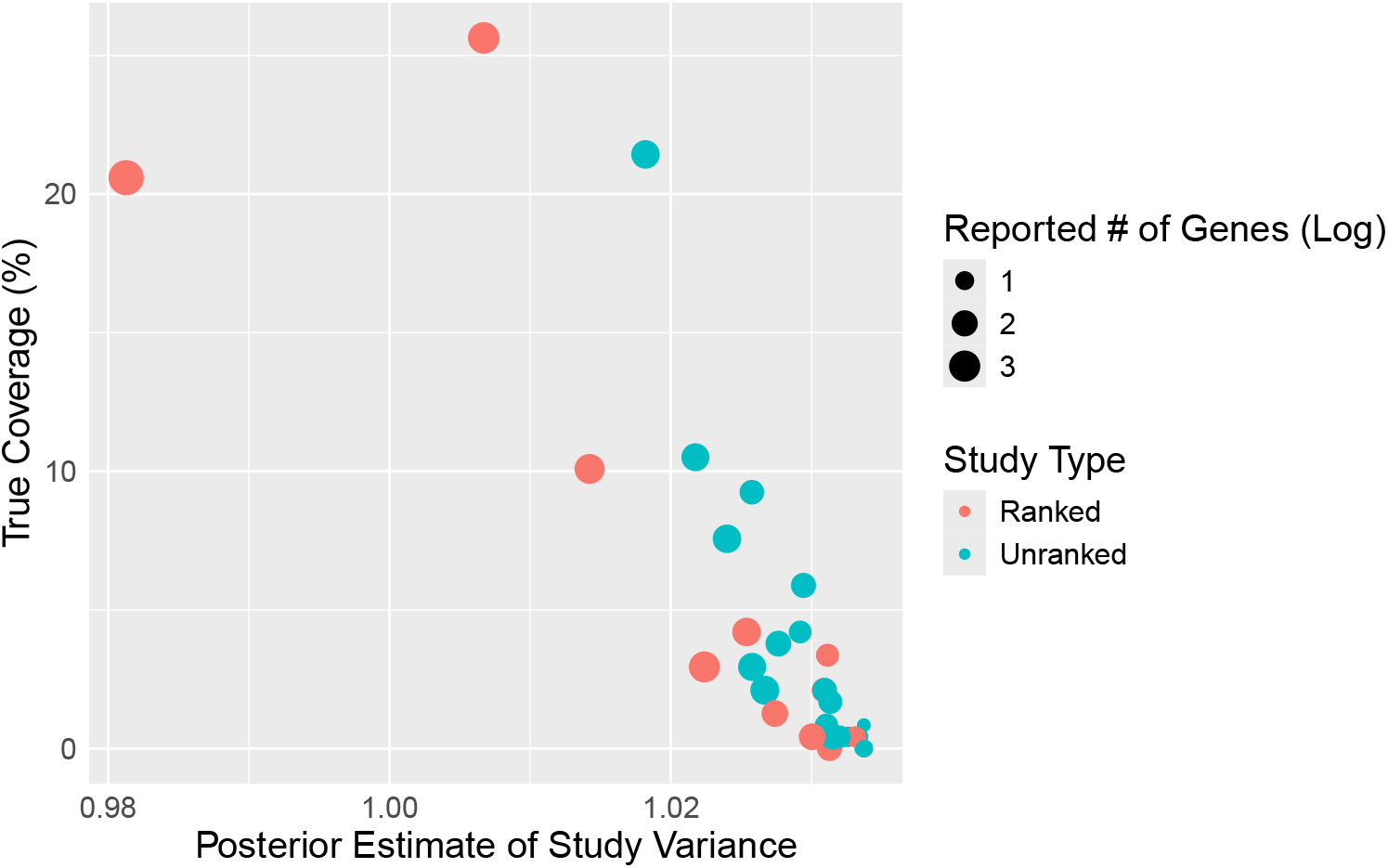
Posterior estimates of study-specific variances in the SARS-COV2 dataset. The posterior estimates of each study in the SARS-COV2 cohort plotted against the true coverage. Dot size indicates the number of reported genes in each study. Color represents the study type: whether they are ranked or unranked.

**Sup. Fig. 4.**
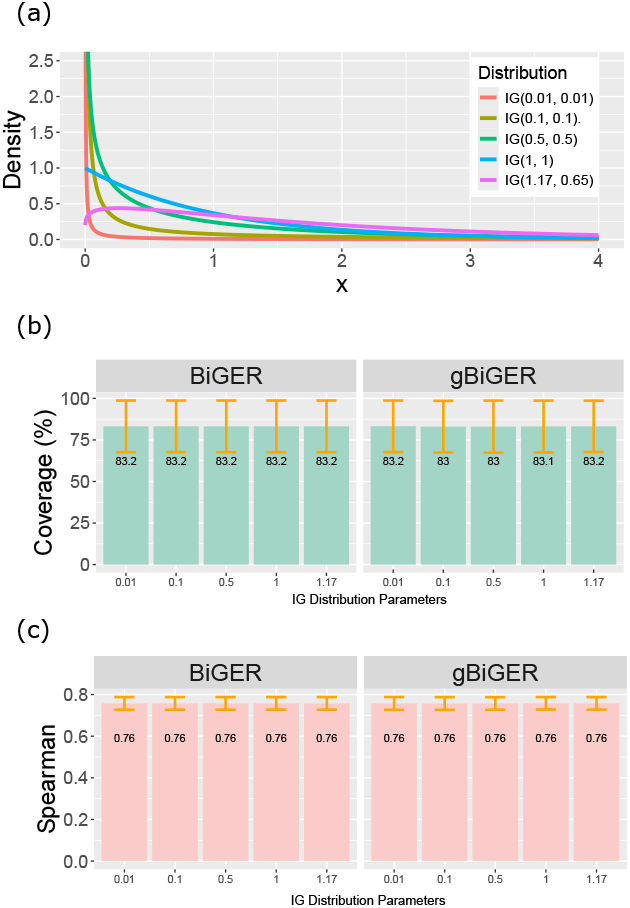
Parameter tuning of Inverse Gamma prior distribution. (a) A comparison of densities of Inverse Gamma distribution with five different distributions. (b-c) The coverage and Spearman’s correlation of the BiGER variants as benchmarked using the default simulation setting. The X-axis denotes the parameters of the Inverse Gamma distribution, where a single value denotes *a* = *b* and two values indicate *a* and *b* respectively. Each bar represents the mean across all 25 replicates, and the orange whiskers denote one standard deviation from the mean. Coverage is presented as percentage in (b).

**Sup. Table 1 Characteristics of three real benchmark datasets**. The table includes each dataset’s gene list type, number of ranked and unranked genes, and the coverage of reported genes. Coverage is calculated using roP genes from each given study.

**Sup. Table 2 Combination of gene types and lists modeled by BiGER**. The table contains a list of all 16 possible combinations with four different gene types. Each combination’s validity, type of list, and additional comments are included.

## Notes

### Competing Interest Statement

The authors have declared no competing interest.

