## Supplementary Note 1 for "BiGER: Bayesian Rank Aggregation in Genomics with Extended Ranking Schemes"

### Supplementary Note 1: Technical Details on the BiGER Family of Models

#### 1 Full List of Notations

In this first section, we list all the notations used throughout the paper and derivations. As a general note, we use subscripts to index along the axes of matrices or vectors, which in most cases correspond to genes and studies. We use letter superscripts as indices instead of exponents. Numeric superscripts are assumed to be exponents by default. For cases when both superscripts and exponents are needed, appropriate parentheses are used for clarity. We divide this section into two first: the first lays out general notations, whereas the latter specifically focuses on variational parameters and MFVI derivations.

##### 1.1 General Notations

- $G$: Total number of genes in the meta-analysis
- $J$: Total number of studies in the meta-analysis
- $j$: Index for each study with $J$ studies in total
- $\mathcal{G}$: Set of all genes included in the meta-analysis
- $\mathcal{G}_j$: Index set of genes in study $j$
- $\mathcal{T}_j^r$: Set of top $n_j^r$ ranked genes in study $j$
- $\mathcal{T}_j^u$: Set of top $n_j^u$ unranked genes provided in study $j$
- $\mathcal{B}_j$: Set of bottom ties in study $j$
- $n_j^r$: The number of top-ranked genes in study $j$ (*i.e.* $|\mathcal{T}_j^r|$)
- $n_j^u$: The number of top-unranked genes in study $j$ (*i.e.* $|\mathcal{T}_j^u|$)
- $n_j^b$: The number of bottom-ties in study $j$ (*i.e.* $|\mathcal{B}_j|$)

- $n_j$: The number of non-missing genes in study $j$.
- $r_{gj}$: The observed rank for gene $g$ in study $j$
- $y^u_{gj}$: The upper bound of $\omega_{gj}$ computed from the rank of gene $g$ in study $j$
- $y^l_{gj}$: The lower bound of $\omega_{gj}$ computed from the rank of gene $g$ in study $j$
- $(r, j)$: The function that returns the index of the gene with rank $r$ in study $j$
- $\boldsymbol{\Theta}$: The collection of all parameters in the BiGER model
- $\omega_{gj}$: The latent weight for gene $g$ in study $j$ (larger $\omega_{gj}$ means smaller $r_{gj}$)
- $\mu_g$: The global importance of gene $g$
- $\epsilon_{gj}$: Random error for gene $g$ in study $j$
- $\sigma^2_j$: Variance for study $j$
- $\mathrm{I}(\cdot)$: Indicator function
- $N(\cdot)$: The Normal distribution with mean-variance parametrization
- $TN(\cdot)$: The Truncated Normal distribution with mean-variance parameterization
- $IG(\cdot)$: The Inverse Gamma distributions with shape-rate parametrization

#### 1.2 Variational Parameters and Derivations

We list all the new notations introduced as part of the VI derivations. In general, we use Greek letters for regular parameters, whereas English letters are used for variational parameters. We use superscripts to indicate the affiliation of each mean and variance parameters.

- $q(\cdot)$: Variational distributions
- $m^\mu_g$: The variational mean parameter for $\mu_g$
- $(s^\mu_g)^2$: The variational variance parameter for $\mu_g$
- $a^\sigma_j$: The variational shape parameter for $\sigma^2_j$
- $b^\sigma_j$: The variational rate parameter for $\sigma^2_j$
- $m^\omega_g$: The variational mean parameter for $\omega_g$

- $(s_g^\omega)^2$: The variational variance parameter for $\omega_g$
- $k$: Index for parameters in the model
- $\mathbb{E}_{-\boldsymbol{\Theta}_k}$: Expectation with respect to all parameters except for $\boldsymbol{\Theta}_k$
- $\Phi(\cdot)$: Standard Normal cumulative density function
- $\varphi(\cdot)$ Standard Normal probability density function

### 2 Variational Inference

First, we write down the full likelihood in its expanded form, and while doing so, we take the logarithm as it is needed in the derivations to follow:

$$\begin{aligned}
\log p(\boldsymbol{y}, \Theta) &= \sum_{j=1}^{J} \sum_{g \in \mathcal{G}_j} \log I(y_{gj}^u < \omega_{gj} < y_{gj}^l) + \sum_{j=1}^{J} \sum_{g \in \mathcal{G}_j} \log N(\omega_{gj} | \mu_g, \sigma_j^2) \\
&\quad + \sum_{g=1}^{G} \log N(\mu_g | 0, 1) + \sum_{j=1}^{J} \log IG(\sigma_j^2 | \alpha, \beta) \\
&= \sum_{j=1}^{J} \sum_{g \in \mathcal{G}_j} \left\{ \log I(y_{gj}^l \leq \omega_{gj} \leq y_{gj}^u) - \log(\sigma_j) + \frac{(\omega_{gj} - \mu_g)^2}{-2\sigma_j^2} \right\} \\
&\quad + \sum_{g=1}^{G} \frac{\mu_g^2}{-2} + \sum_{j=1}^{J} (-\alpha - 1) \log(\sigma_j^2) - \frac{\beta}{\sigma_j^2} + c\,.
\end{aligned}$$

Note here that we use $c$ as a generic normalizing constant without distinguishing between their specific values. Again, for all expectations in the following derivations, they are taken with respect to their enclosed parameters unless otherwise notated.

#### 2.1 Variational Update for $\mu_g$

We update each $\mu_g$ individually:

$$\begin{aligned}
q(\mu_g) &\propto \mathbb{E}_{-\mu_g} \log p\left(\mu_g | \boldsymbol{y}, \Theta_{-\mu_g}\right) \\
&= \sum_{j: g \in \mathcal{G}_j} \left\{ \mathbb{E}_\omega \mathbb{E}_{\sigma^2} \frac{(\omega_{gj} - \mu_g)^2}{-2\sigma_j^2} \right\} + \frac{\mu_g^2}{-2} + c \\
&= \sum_{j: g \in \mathcal{G}_j} \left\{ \mathbb{E}_\omega (\omega_{gj} - \mu_g)^2 \times \left(\frac{1}{-2}\right) \mathbb{E}_{\sigma^2} \left(\frac{1}{\sigma_j^2}\right) \right\} + \frac{\mu_g^2}{-2} + c
\end{aligned}$$

$$
\begin{aligned}
&= \sum_{j:g\in\mathcal{G}_j} \left\{ \left(\frac{1}{-2}\right) \mathbb{E}\left(\frac{1}{\sigma_j^2}\right) \left[\mathbb{E}(\omega_{gj}^2) - 2\mathbb{E}(\omega_{gj})\mu_g + \mu_g^2\right] \right\} + \frac{\mu_g^2}{-2} + c \\
&= \mu_g \sum_{j:g\in\mathcal{G}_j} \left\{ \mathbb{E}\left(\frac{1}{\sigma_j^2}\right) \mathbb{E}(\omega_{gj}) \right\} - \mu_g^2 \left[ \sum_{j:g\in\mathcal{G}_j} \frac{1}{2}\mathbb{E}\left(\frac{1}{\sigma_j^2}\right) + \frac{1}{2} \right] + c\,.
\end{aligned}
$$

This is obvious a quadratic form, meaning that it is a normal distribution with the following specification

$$
q(\mu_g) \sim N\left( m_g^\mu = \frac{\sum_{j:g\in\mathcal{G}_j} \left\{ \mathbb{E}\left(\frac{1}{\sigma_j^2}\right) \mathbb{E}(\omega_{gj}) \right\}}{\sum_{j:g\in\mathcal{G}_j} \mathbb{E}\left(\frac{1}{\sigma_j^2}\right) + 1}, (s_g^\mu)^2 = \frac{1}{\sum_{j:g\in\mathcal{G}_j} \mathbb{E}\left(\frac{1}{\sigma_j^2}\right) + 1} \right)\,.
$$

As expected, this formulation is very similar to that of a Gibbs simpler used in BiGER.

#### 2.2 Variational Update for $\sigma_j^2$

For the study-specific variance, each $\sigma_j^2$ is dealt with individually as well:

$$
\begin{aligned}
q(\sigma_j^2) &\propto \mathbb{E}_{-\sigma_j^2} \log p\left(\sigma_j^2 | \boldsymbol{y}, \Theta_{-\sigma_j^2}\right) \\
&= \sum_{g\in\mathcal{G}_j} -\frac{1}{2}\log\sigma_j^2 - \mathbb{E}_{-\sigma_j^2}\left[\frac{(\omega_{gj}-\mu_g)^2}{2\sigma_j^2}\right] + (-\alpha-1)\log(\sigma_j^2) - \frac{\beta}{\sigma_j^2} + c \\
&= -\frac{n_j}{2}\log\sigma_j^2 - \sum_{g\in\mathcal{G}_j} \frac{\mathbb{E}(\omega_{gj}^2) - 2\mathbb{E}(\omega_{gj})\mathbb{E}(\mu_g) + \mathbb{E}(\mu_g^2)}{2\sigma_j^2} + (-\alpha-1)\log\sigma_j^2 - \frac{\beta}{\sigma_j^2} + c \\
&= -(\frac{1}{2}n_j + \alpha + 1)\log\sigma_j^2 - \frac{2\beta + \sum_{g\in\mathcal{G}_j} \mathbb{E}(\omega_{gj}^2) - 2\mathbb{E}(\omega_{gj})\mathbb{E}(\mu_g) + \mathbb{E}(\mu_g^2)}{2\sigma_j^2} + c\,.
\end{aligned}
$$

Again, this is a Inverse-Gamma distribution just as expected:

$$
q(\sigma_j^2) \sim \text{Inv-Gamma}\left( a_j^\sigma = \frac{1}{2}n_j + \alpha, b_j^\sigma = \beta + \sum_{g\in\mathcal{G}_j} \frac{1}{2}\mathbb{E}(\omega_{gj}^2) - \mathbb{E}(\omega_{gj})\mathbb{E}(\mu_g) + \frac{1}{2}\mathbb{E}(\mu_g^2) \right)
$$

#### 2.3 Variational Update for $\omega_{gj}$

Finally, we need to deal with the weights for each gene in each study. We have

$$
\begin{aligned}
q(\omega_{gj}) &\propto \mathbb{E}_{-\omega_{gj}} \log p\left(\omega_{gj} | \boldsymbol{y}, \Theta_{-\omega_{gj}}\right) \\
&= \log I(y_{gj}^l \le \omega_{gj} \le y_{gj}^u) + \mathbb{E}_{-\omega_{gj}}\left[\frac{(\omega_{gj}-\mu_g)^2}{-2\sigma_j^2}\right] + c
\end{aligned}
$$

$$= \log I(y_{gj}^l \leq \omega_{gj} \leq y_{gj}^u) + \mathbb{E}\left(\frac{1}{-2\sigma_j^2}\right)\left(\omega_{gj}^2 - 2\mathbb{E}(\mu_g)\omega_{gj} + \mathbb{E}(\mu_g^2)\right) + c$$

$$= \log I(y_{gj}^l \leq \omega_{gj} \leq y_{gj}^u) + \frac{(\omega_{gj} - \mathbb{E}(\mu_g))^2}{-2\mathbb{E}\left(\frac{1}{\sigma_j^2}\right)^{-1}} + c\,,$$

which is another truncated normal distribution. The distribution is

$$q(\omega_{gj}) \sim TN\left(m_{gj}^\omega = \mathbb{E}(\mu_g), (s_{gj}^\omega)^2 = \mathbb{E}\left(\frac{1}{\sigma_j^2}\right)^{-1}, y_{gj}^l, y_{gj}^u\right)$$

with $y_{gj}^l$ and $y_{gj}^u$ as truncation bounds.

#### 2.4 Detour: Why not the original parametrization?

Here, we take a slight detour to show that under the original BiGER model, the boundaries are problematic with expectations. Under the BiGER parametrization, it is easy to show that

$$\begin{aligned} q(\omega_{gj}) &\propto \mathbb{E}_{-\omega_{gj}} \log p(\omega_{gj}|\boldsymbol{y}, \Theta_{-\omega_{gj}}) \\ &= \mathbb{E}_{-\omega_{gj}}\left\{\mathrm{I}\left(\omega_{(1,j),j} > \omega_{(2,j),j} > \ldots > \omega_{(n_j^r,j),j} > \sup_{g_u \in \mathcal{T}_j^u} \omega_{g_u j} \geq \inf_{g_u \in \mathcal{T}_j^u} \omega_{g_u j} > \sup_{g_b \in \mathcal{B}_j} \omega_{g_b j}\right)\right\} \\ &\quad + \mathbb{E}_{-\omega_{gj}}\left[\frac{(\omega_{gj} - \mu_g)^2}{-2\sigma_j^2}\right] + c \\ &= \mathbb{E}_{-\omega_{gj}} \mathrm{I}\left(\omega_{gj}^l < \omega_{gj} < \omega_{gj}^u\right) + \mathbb{E}_{-\omega_{gj}}\left[\frac{(\omega_{gj} - \mu_g)^2}{-2\sigma_j^2}\right] + c\,. \end{aligned}$$

Without loss of generality, we assume that $\omega_{gj}$ is a top-ranked gene with genes ranked both above and below it. In a slight abuse of notation for convenience, we use the notations $\omega_{gj}^l$ and $\omega_{gj}^u$ to denote the upper and lower bounds of $\omega_{gj}$. Note that the last line follows from the fact that only the upper and lower bound for $\omega_{gj}$ matter. It is obvious that in this parametrization, the bounds depend on other latent weights $w$, which are also parameters. Therefore, we cannot easily take or distribute the double expectations, and we do not have a TN distribution in closed form.

#### 2.5 Expectations

In the variational updates, we need to find the expectations regarding the variational distributions. They are:

$$
\begin{aligned}
\mathbb{E}(\mu_g) &= m_g^\mu \\
\mathbb{E}(\mu_g^2) &= \text{var}(\mu_g) + \mathbb{E}(\mu_g)^2 = (s_g^\mu)^2 + (m_g^\mu)^2 \\
\mathbb{E}\left(\frac{1}{\sigma_j^2}\right) &= \frac{a_j^\sigma}{b_j^\sigma} \\
\mathbb{E}(\omega_{gj}) &= m_{gj}^\omega + \frac{\varphi(y_{gj}^l) - \varphi(y_{gj}^u)}{\Phi(y_{gj}^u) - \Phi(y_{gj}^l)} s_{gj}^\omega \\
\mathbb{E}(\omega_{gj}^2) &= \left[m_{gj}^\omega + \frac{\varphi(y_{gj}^l) - \varphi(y_{gj}^u)}{\Phi(y_{gj}^u) - \Phi(y_{gj}^l)} s_{gj}^\omega\right]^2 \\
&\quad + (s_{gj}^\omega)^2 \left[1 - \frac{y_{gj}^u \varphi(y_{gj}^u) - y_{gj}^l \varphi(y_{gj}^l)}{\Phi(y_{gj}^u) - \Phi(y_{gj}^l)} - \left(\frac{\varphi(y_{gj}^l) - \varphi(y_{gj}^u)}{\Phi(y_{gj}^u) - \Phi(y_{gj}^l)}\right)^2\right]
\end{aligned}
$$

As a note of optimization, we store the parameters in each iteration rather than re-computing everything on the fly. Further, $\mathbb{E}(\omega_{gj})$ can be calculated from existing packages without the need for reimplementation.

#### 2.6 The ELBO

Typically, the evidence lower bound (ELBO) is used to monitor convergence, and we derive it for completeness. The ELBO from MFVI is defined as

$$
\begin{aligned}
\text{ELBO}(q) &= \mathbb{E}[\log p(\Theta)] + \mathbb{E}[\log p(\boldsymbol{y}|\Theta)] - \mathbb{E}[\log q(\Theta)] \\
&= \mathbb{E}[\log p(\Theta)] - \mathbb{E}[\log q(\Theta)] \,.
\end{aligned}
$$

Note here that the term $\mathbb{E}[\log p(\boldsymbol{y}|\Theta)]$ disappears because $\log p(\boldsymbol{y}|\Theta)$ is an indicator, and in our algorithm, we ensure that the the ordering is always correct through the truncated normal distribution. Now, let's proceed:

$$
\begin{aligned}
\text{ELBO}(q) &= \mathbb{E}(\log p(\boldsymbol{\omega})) + \mathbb{E}(\log p(\boldsymbol{\mu})) + \mathbb{E}(\log p(\boldsymbol{\sigma}^2)) - \mathbb{E}[\log q(\boldsymbol{\omega})] - \mathbb{E}[\log q(\boldsymbol{\mu})] - \mathbb{E}[\log q(\boldsymbol{\sigma^2})] \\
&= \sum_{j=1}^{J} \sum_{g \in \mathcal{G}_j} \mathbb{E}[\log p(\omega_{gj})] + \sum_{g=1}^{G} \mathbb{E}(\log p(\mu_g)) + \sum_{j=1}^{J} \mathbb{E}(\log p(\sigma_j^2)) - \sum_{j=1}^{J} \sum_{g \in \mathcal{G}_j} \mathbb{E}[\log q(\omega_{gj})] \\
&\quad - \sum_{g=1}^{G} \mathbb{E}[\log q(\mu_g)] - \sum_{j=1}^{J} \mathbb{E}[\log q(\sigma_j^2)]
\end{aligned}
$$

$$
\begin{aligned}
&= \sum_{j=1}^{J}\sum_{g\in\mathcal{G}_j}\left\{-\mathbb{E}(\log(\sigma_j)) + \mathbb{E}\left[\frac{(\omega_{gj}-\mu_g)^2}{-2\sigma_j^2}\right]\right\} + \sum_{g=1}^{G}\frac{\mathbb{E}(\mu_g^2)}{-2} + \sum_{j=1}^{J}(-\alpha-1)\mathbb{E}(\log(\sigma_j^2)) \\
&\quad - \mathbb{E}\left(\frac{\beta}{\sigma_j^2}\right) - \sum_{j=1}^{J}\sum_{g\in\mathcal{G}_j}\left\{-\log s_{gj}^{\omega} + \mathbb{E}\left[\frac{(\omega_{gj}-m_{gj}^{\omega})^2}{-2(s_{gj}^{\omega})^2}\right]\right\} \\
&\quad - \sum_{g=1}^{G}\left\{-\log(s_g^{\mu}) + \mathbb{E}\left[\frac{(\mu_g-m_g^{\mu})^2}{-2(s_g^{\mu})^2}\right]\right\} - \sum_{j=1}^{J}\left\{(-a_j^{\sigma}-1)\mathbb{E}(\log(\sigma_j^2)) - \mathbb{E}\left(\frac{b_j^{\sigma}}{\sigma_j^2}\right)\right\} + c \\
&= \sum_{j=1}^{J}\sum_{g\in\mathcal{G}_j}\left\{-\mathbb{E}(\log(\sigma_j)) + \mathbb{E}\left[\frac{(\omega_{gj}-\mu_g)^2}{-2\sigma_j^2}\right]\right\} + \sum_{g=1}^{G}\frac{(m_g^{\mu})^2+(s_g^{\mu})^2}{-2} \\
&\quad + \sum_{j=1}^{J}(-\alpha-1)\mathbb{E}(\log(\sigma_j^2)) - \frac{\beta a_j^{\sigma}}{b_j^{\sigma}} - \sum_{j=1}^{J}\sum_{g\in\mathcal{G}_j}\left\{-\log s_{gj}^{\omega} + \mathbb{E}\left[\frac{(\omega_{gj}-m_{gj}^{\omega})^2}{-2(s_{gj}^{\omega})^2}\right]\right\} \\
&\quad - \sum_{g=1}^{G}\left\{-\log(s_g^{\mu}) + \mathbb{E}\left[\frac{(\mu_g-m_g^{\mu})^2}{-2(s_g^{\mu})^2}\right]\right\} - \sum_{j=1}^{J}\left\{(-a_j^{\sigma}-1)\mathbb{E}(\log(\sigma_j^2)) - a_j^{\sigma}\right\} + c
\end{aligned}
$$

We do not compute this in our algorithm for efficiency reasons as there is no closed form for some of the expectations in the ELBO. In our algorithm, we monitored the change in parameters for convergence as noted in the **Methods** section.
